# Control of mitochondrial dynamics by dPGC1 limits Yorkie-induced oncogenic growth in *Drosophila*

**DOI:** 10.1101/2024.12.02.626378

**Authors:** Wei Qi Guinevere Sew, Maria Molano-Fernández, Zhiquan Li, Artim Lange, Nahia Pérez de Ciriza, Lene Juel Rasmussen, Héctor Herranz

**Affiliations:** Department of Cellular and Molecular Medicine, University of Copenhagen, 2200 Copenhagen N, Denmark; Centre for Healthy Aging, Department of Cellular and Molecular Medicine, University of Copenhagen, 2200 Copenhagen N, Denmark

**Keywords:** mitochondria, mitochondrial dynamics, *Drosophila*, Hippo pathway, cancer, cell cycle

## Abstract

Mitochondrial function and dynamics are essential for maintaining cellular homeostasis and overall health. Disruptions in these processes can contribute to various diseases, including cancer. The Hippo signaling pathway, a key regulator of tissue growth, plays a central role in cancer through its main effector, the Yes-associated protein (YAP), known as Yorkie (Yki) in *Drosophila*. In this model organism, *Yki* upregulation drives benign tissue overgrowth in imaginal discs. Our research shows that the conserved metabolic regulator dPGC1 restricts Yki-driven tissue hyperplasia and helps maintain epithelial integrity *in vivo*. Combined *Yki* upregulation and *dPGC1* depletion results in tumors characterized by enlarged mitochondria and upregulation of genes promoting mitochondrial fusion, a condition that is both necessary and sufficient for Yki-driven oncogenic growth. We further demonstrate that mitochondrial enlargement is associated with increased levels of the cell cycle regulator Cyclin E, which is critical for tumor development. These findings identify dPGC1 as a context-dependent tumor suppressor that coordinates mitochondrial dynamics and cell cycle regulation in response to oncogene activation, with implications for understanding cancer development in humans.

## Introduction

Cancer is a multistep process that involves the accumulation of mutations affecting different cellular functions known as the hallmarks of cancer. These include central cellular processes such as cell proliferation, apoptosis, and cellular energetics. Together, these hallmarks promote the transformation of normal cells into malignant entities (Hanahan, 2022; Hanahan & Weinberg, 2011). Tumors frequently undergo major changes in their metabolism, which is required to fulfill the high energetic demands of cancer cells. Mitochondria are cytoplasmic organelles that play central roles in cellular energetics. Alterations in mitochondrial function can provide plasticity and adaptability to tumor cells growing in harsh environments, such as hypoxia, reduced nutrient availability, and exposure to cancer treatments (Vyas *et al*, 2016). Consequently, mitochondrial alterations influence various aspects of tumor initiation and disease progression, and genetic changes affecting mitochondria can increase cancer risk (Pavlova & Thompson, 2016; Sullivan *et al*, 2016).

Mitochondria are highly dynamic organelles. Continuous fusion and fission events are crucial for maintaining mitochondrial integrity and function. Mitochondrial fission mediates the division of one mitochondrion into two and is essential for processes such as mitosis and quality control. Mitochondrial fusion, by contrast, merges two mitochondria into one, allowing the exchange of mitochondrial contents and supporting function, especially under cellular stress (Adebayo *et al*, 2021; Mishra & Chan, 2014). Beyond maintaining mitochondrial integrity and function, the balance between fission and fusion is crucial for regulating the cell cycle (Antico Arciuch *et al*, 2012; Qian *et al*, 2012). The cell cycle ensures that a cell replicates and segregates its genome faithfully before it divides into two identical daughter cells. Errors in this process could lead to genomic instability, which is a cancer driver (Tubbs & Nussenzweig, 2017). Defective mitochondrial dynamics can result in an imbalance between fusion and fission, which has been associated with human diseases, including cancer. Therefore, mitochondrial dynamics must be tightly regulated to ensure proper cellular function and tissue homeostasis (Archer, 2013; Dai & Jiang, 2019; Liao *et al*, 2024; Picard *et al*, 2013; Senft & Ronai, 2016; Yu *et al*, 2023).

The Peroxisome Proliferator Activated Receptor Gamma Coactivator-1 (PGC1) family of transcriptional regulators plays central roles in controlling mitochondrial biogenesis and function. This family was originally characterized by the Spiegelman lab, who identified PGC1α as a cold-inducible coactivator of nuclear receptors involved in adaptive thermogenesis and mitochondrial regulation (Puigserver *et al*, 1998). Subsequent work from the same group demonstrated that PGC1α orchestrates mitochondrial gene expression and respiration (Wu *et al*, 1999), regulates oxidative metabolism and muscle fiber type specification (Lin *et al*, 2002), and promotes mitochondrial biogenesis and oxidative phosphorylation in skeletal muscle cells (Lin *et al*, 2004). In mammals, this family includes PGC1α, PGC1β, and PGC1-related coactivator (PRC) (Scarpulla, 2011; Villena, 2015). These family members share structural and functional similarities. Among them, PGC1α has been most extensively studied in the context of cancer, where it regulates mitochondrial dynamics, metabolic reprogramming, and tumor progression. In contrast, PGC1β and PRC have primarily been characterized in normal physiology, where they regulate mitochondrial biogenesis. While PGC1β and PRC are primarily associated with cellular metabolism and energy homeostasis, PGC1β also contributes to mitochondrial fusion, notably through the regulation of MFN2 and other mitochondrial factors (Liesa *et al*, 2008).

PGC1α controls mitochondrial dynamics by regulating the expression of genes involved in fusion and fission, ensuring a proper balance between these processes to preserve mitochondrial integrity (Abu Shelbayeh *et al*, 2023; Chen *et al*, 2022). The role of PGC1α in cancer is complex and context dependent. It can act both as both a tumor suppressor and as an oncogenic factor, depending on the cancer type and cellular context (Bhalla *et al*, 2011; Girnun, 2012; Jones *et al*, 2012; Tan *et al*, 2016; Torrano *et al*, 2016; Wang *et al*, 2024). Therefore, elucidating the context-specific functions of PGC1α in cancer is crucial for developing targeted therapies. In cancers where PGC1α acts as a tumor suppressor, strategies to enhance its activity might be beneficial. In contrast, in cancers where PGC1α supports tumor growth, inhibiting its function could be a potential therapeutic approach.

*Drosophila* is emerging as a valuable *in vivo* model to study various aspects of cancer (Gonzalez, 2013). Notably, *Drosophila* possesses a sole *PGC1* orthologue: *dPGC1*, also known as *spargel*, which has also been involved in the regulation of energy metabolism and mitochondrial function (Gershman *et al*, 2007; Mukherjee *et al*, 2014). The functional homology between dPGC1 and mammalian PGC1 members facilitates the study of PGC1 functions in *Drosophila* without the complications of redundancy.

The Hippo signaling pathway is a conserved regulator of normal and oncogenic growth. The central elements of this pathway comprise a regulatory kinase cascade that, when active, limits tissue growth by phosphorylating and inhibiting the activity of transcriptional coactivator with PDZ-binding motif (TAZ) and Yes-associated protein (YAP), known as Yorkie (Yki) in *Drosophila* (Zheng & Pan, 2019). Deregulation of the activity of the Hippo pathway affects tumor development and has been associated with traits of oncogenesis such as cell proliferation and survival, cancer metabolism, invasion, and metastasis (Barron & Kagey, 2014; Moon *et al*, 2018). Consistent with their growth-regulatory roles, YAP and TAZ are frequently amplified, while upstream elements in the pathway are mutated in different cancer types (Calses *et al*, 2019).

Work from Banerjee’s group (Nagaraj *et al*, 2012) demonstrated that activation of the Hippo pathway effector Yki in *Drosophila* leads to increased mitochondrial fusion through the transcriptional upregulation of key fusion genes such as *Mitofusin* (*dMfn* also known as *Marf*) and *Optic atrophy 1* (*Opa1*). Their study showed that *Yki* overexpression alone is sufficient to induce mitochondrial elongation and to upregulate several mitochondrial genes, including those involved in fusion and oxidative stress responses, while notably not affecting *dPGC1* expression. These findings suggest that Yki can directly modulate mitochondrial structure and function independently of dPGC1, and that the regulation of mitochondrial fusion genes contributes to the growth-promoting role of Yki in epithelial tissues. Building on this foundation, our study explores how *dPGC1* depletion further modulates the transcriptional and morphological mitochondrial responses in a *Yki*-overexpressing context. By comparing *Yki* overexpression alone to *Yki* upregulation combined with *dPGC1* knockdown, we dissect the specific contributions of dPGC1 not only to mitochondrial dynamics and gene regulation, but also to the tumor-like growth properties driven by Yki activation.

We find that while depletion of the transcriptional coactivator *dPGC1* has a minor impact on normal *Drosophila* wing imaginal disc development, only causing a subtle growth defect, it dramatically drives tumor growth and cellular transformation in imaginal discs overexpressing *Yki*. This establishes dPGC1 as a context-dependent tumor suppressor that specifically limits Yki-driven tissue overgrowth without altering *Yki* expression or activity, and notably, not affecting overgrowth by other oncogenes. Tumors driven by *Yki* upregulation and *dPGC1* depletion exhibit enlarged mitochondria, a phenotype that correlates with the transcriptional upregulation of key mitochondrial fusion genes, *dMfn* and *Opa1*. We demonstrate that this is both necessary and sufficient to promote Yki-driven oncogenic growth and malignancy. Tumors driven by *Yki* overexpression and *dPGC1* knockdown show increased levels of Cyclin E protein. Cyclin E is essential for tumor development in this context, promoting growth and inducing DNA damage, two hallmarks of cancer. This study elucidates a novel connection between aberrant mitochondrial dynamics, defective cell cycle regulation, and DNA damage in oncogenic processes occurring *in vivo*, suggesting that targeting mitochondrial fusion components may offer new therapeutic strategies to limit Yki/YAP-driven tumor growth.

## Results

### dPGC1 is Required for Normal Wing Growth

The systemic functions of dPGC1 have been extensively studied, and it has been shown to control central physiological processes such as early development, tissue homeostasis, obesity, and aging (Chen *et al*, 2012; Diop *et al*, 2015; George & Jacobs, 2019; Mukherjee *et al*., 2014; Mukherjee & Duttaroy, 2013; Rera *et al*, 2011; Rogers & Rogina, 2014). However, its role in highly proliferative tissues such as developing organs or neoplastic growth remains poorly understood. To address this gap, we employed the *Drosophila* wing imaginal disc, a well-established *in vivo* model composed of a proliferative epithelial monolayer that gives rise to the adult wing and thorax during metamorphosis. This tissue serves as a powerful system for investigating epithelial development, growth regulation, and tumorigenesis (Herranz *et al*, 2016).

To dissect the function of dPGC1 in this context, we examined the wings of *dPGC1^1^* mutant flies, which carry a P-element insertion in the *dPGC1* gene. This insertion disrupts normal transcription and results in reduced gene function (Tiefenbock *et al*, 2010). Despite this molecular alteration, *dPGC1^1^* homozygous mutants are viable and do not display overt morphological abnormalities. These observations are consistent with studies in mice showing that PGC1α and PGC1β are not essential for normal development (Lelliott *et al*, 2006; Leone *et al*, 2005; Lin *et al*., 2004; Sonoda *et al*, 2007; Vianna *et al*, 2006). We detected a modest but statistically significant reduction in the size of both wing discs and adult wings in *dPGC1^1^/dPGC1^1^*mutants compared to controls (Fig 1A, B; Fig S1). These results indicate that dPGC1, while not essential for viability or gross morphology, plays a role in promoting normal wing growth. To confirm this role, we employed the Gal4-UAS binary system to manipulate *dPGC1* expression in a spatially and temporally controlled manner (Brand & Perrimon, 1993). Using the MS1096-Gal4 driver, which is active in the wing disc epithelium (Fig 1C), we specifically modulated gene activity during wing development and assessed the resulting adult phenotypes. Consistent with the analysis of *dPGC1* mutants, expression of *dPGC1-RNAi* transgenes under the control of MS1096-Gal4 led to a modest size reduction (Fig 1D, E). The efficiency of the transgenes used to deplete *dPGC1* is shown in Fig S2. Overexpression of *dPGC1* did not affect wing size or pattern (Fig 1D, E). These results indicate that, although dispensable for *Drosophila* viability, dPGC1 is required for normal wing growth.

**Figure 1.**
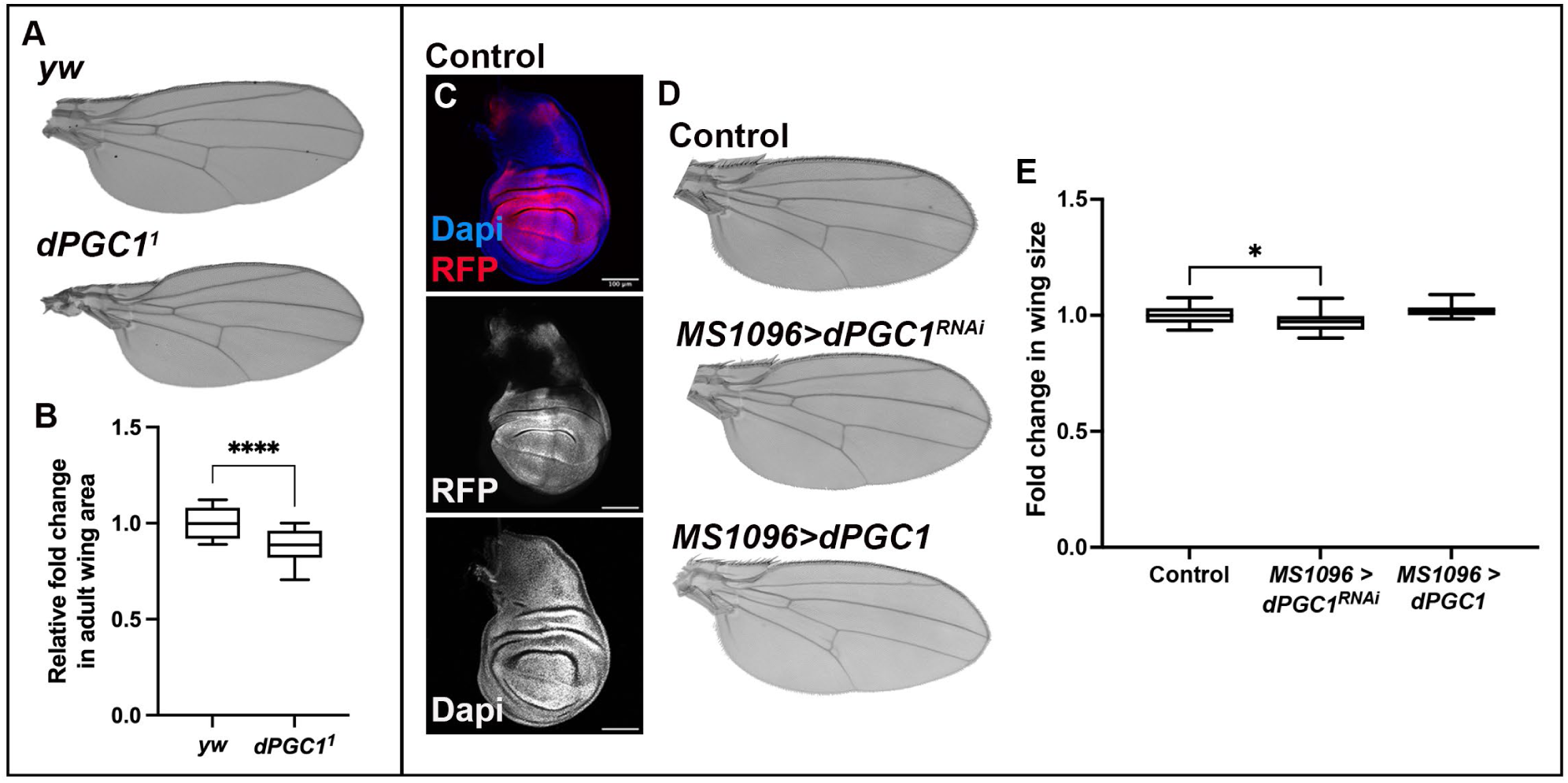
Reduced dPGC1 Activity Leads to Smaller Wings. **(A)** Cuticle preparations of adult wings from the following genotypes: *yw/yw* (control) and *dPGC1^1^/dPGC1^1^* (mutant). **(B)** Quantification of adult wing size of the genotypes indicated in A. Wing area is normalized to the mean of the control (*yw*). Statistical significance was determined using an unpaired t-test (n = 20 [*yw*], n = 20 [*dPGC1^1^*]). ****, p < 0.0001. **(C)** Confocal micrograph showing a *MS1096-Gal4, UAS-RFP* third instar wing imaginal disc. RFP labels the region of the disc expressing this Gal4 line and is shown in red. DAPI labels the DNA and is shown in blue. Scale bar, 100 μm. **(D)** Cuticle preparations of adult wings from *MS1096-Gal4, UAS-LacZ* control flies; *MS1096-Gal4, UAS-dPGC1-RNAi* flies; and *MS1096-Gal4, EP-dPGC1* flies. **(E)** Quantification of adult wing size of the genotypes indicated in D. Wing area is normalized to the mean of the control (*MS1096>LacZ*). Statistical significance was determined using unpaired t-tests with Welch’s correction (n = 20 [*MS1096>LacZ*], n = 20 [*MS1096>dPGC1-RNAi*], n = 20 [*MS1096>dPGC1*]). *p < 0.05.

### dPGC1 Restricts Yki-Driven Tissue Overgrowth in *Drosophila*

Given the role of dPGC1 in supporting normal wing growth, we next sought to determine whether this transcriptional regulator also influences pathological growth conditions. Since many of the molecular mechanisms that govern normal development are co-opted during tumorigenesis, we hypothesized that dPGC1 might modulate the cellular response to oncogenic signals. Among the various oncogenic drivers in *Drosophila*, we started analyzing the Hippo pathway effector Yki, a conserved proto-oncogene that promotes tissue overgrowth and modulates mitochondrial structure and function, processes in which dPGC1 is also implicated (Gerlach *et al*, 2018; Gerlach *et al*, 2019; Huang *et al*, 2005; Nagaraj *et al*., 2012; Nolo *et al*, 2006; Thompson & Cohen, 2006; Tiefenbock *et al*., 2010). This functional overlap in mitochondrial regulation provided a strong rationale to explore whether dPGC1 modulates Yki-driven tumor-like growth. To test this, we depleted *dPGC1* in wing imaginal discs overexpressing *Yki* using the apterous-Gal4 (ap-Gal4) driver, which has been successfully employed to dissect various aspects of tumor formation (Eichenlaub *et al*, 2018; Gerlach *et al*., 2018; Herranz *et al*, 2012; Herranz *et al*, 2014; Song *et al*, 2019). As previously reported, *Yki* overexpression leads to a marked increase in tissue size (Fig 2A, C, E) (Huang *et al*., 2005). While *dPGC1* knockdown alone did not affect wing disc size, its depletion in the context of *Yki* overexpression significantly enhanced the overgrowth phenotype (Fig 2A–E). These findings indicate that, although dPGC1 contributes modestly to normal wing development, it plays a critical role in restraining Yki-driven tissue expansion.

**Figure 2.**
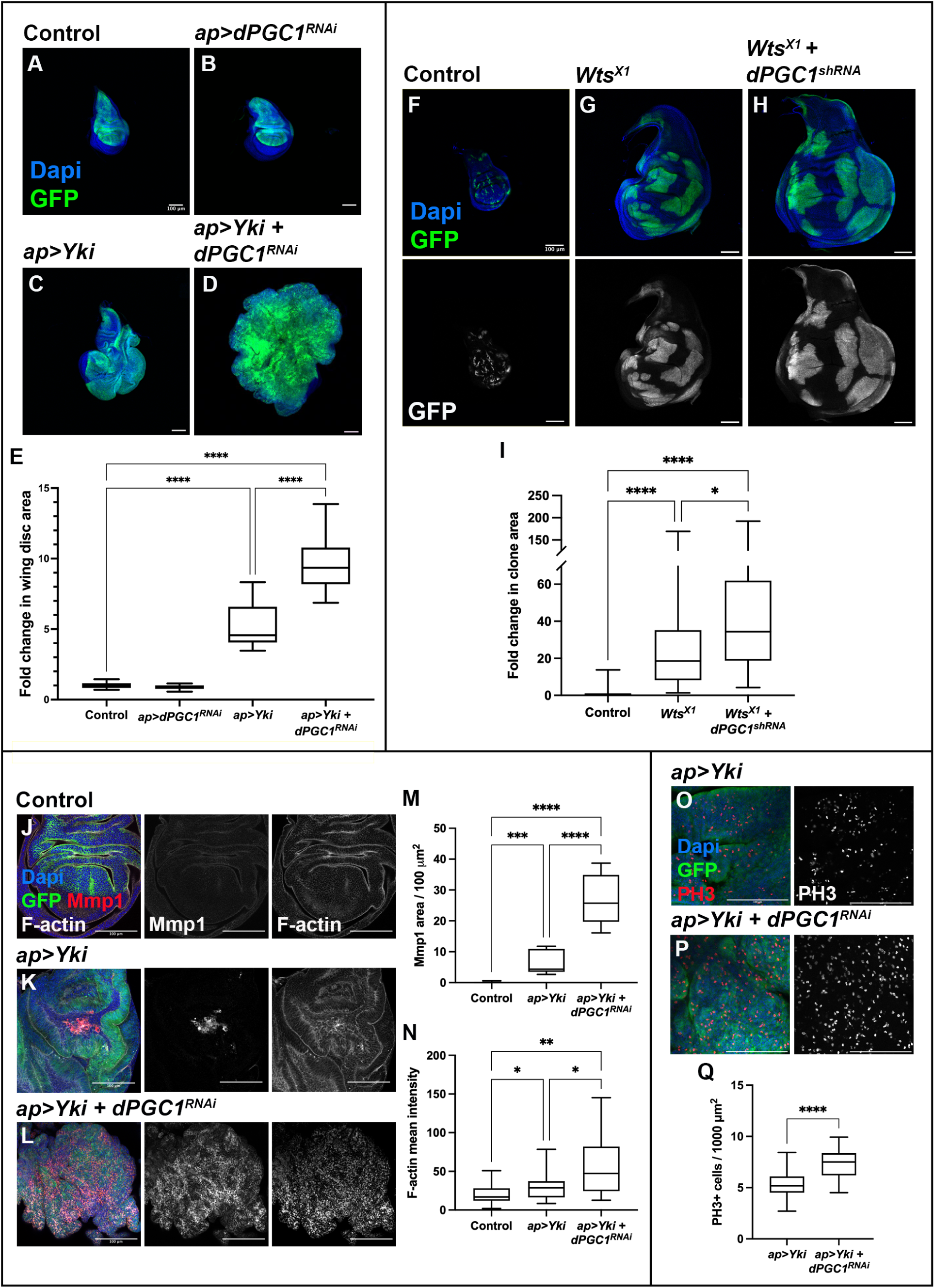
Oncogenic Growth in Discs Overexpressing *Yki* and Downregulating *dPGC1*. **(A-D)** Confocal micrographs showing discs of the following genotypes: *ap-Gal4, UAS-GFP, UAS-LacZ* (A); *ap-Gal4, UAS-GFP, UAS-dPGC1-RNAi* (B); *ap-Gal4, UAS-GFP, UAS-Yki, UAS-LacZ* (C); and *ap-Gal4, UAS-GFP, UAS-Yki, UAS-dPGC1-RNAi* (D). GFP is shown in green, and DAPI labels the DNA and is shown in blue. Scale bars, 100 μm. **(E)** Quantification of GFP-positive area in third instar wing imaginal discs of the genotypes shown in A-D. GFP-positive area is normalized to the mean of the control *(ap>LacZ*). Statistical significance was determined using unpaired t-tests with Welch’s correction (n = 18 [*ap>LacZ*], n = 19 [*ap>dPGC1-RNAi*], n = 20 [*ap>Yki*], n = 20 *[ap>Yki, dPGC1-RNAi*]). ****p < 0.0001. **(F-H)** Confocal micrographs showing representative images of imaginal discs with GFP-labeled MARCM control clones (F), *Wts^X1^* clones (G), and *Wts^X1^* clones expressing *UAS-dPGC1-shRNA* (H). GFP is shown in green, and DAPI labels the DNA and is shown in blue. Scale bars, 100 μm. **(I)** Quantification of GFP-positive clones in the genotypes shown in F-H. GFP-positive area is normalized to the mean of the control. Statistical significance was determined using a Kruskal Wallis test for non-parametric data (n = 102 [Control], n = 92 [*Wts^X1^*], n = 65 [*Wts^X1^ + dPGC1^shRNA^*]). *p < 0.05, ****p < 0.0001. **(J-L)** Confocal micrographs of imaginal wing discs of the following genotypes: *ap-Gal4, UAS-GFP, UAS-LacZ* (J); *ap-Gal4, UAS-GFP, UAS-Yki, UAS-LacZ* (K); and *ap-Gal4, UAS-GFP, UAS-Yki, UAS-dPGC1-RNAi* (L). F-actin labels cell polarity and is shown in grayscale. Mmp1 is shown in red. GFP is shown in green. DAPI labels the DNA and is shown in blue. Scale bars, 100 μm. (**M**) Quantification of Mmp1 area per GFP-positive area in the genotypes shown in J-L. Statistical significance was determined using a Brown-Forsythe and Welch ANOVA test (n = 12 [*ap>LacZ*], n = 11 [*ap>Yki*], n = 11 *[ap>Yki, dPGC1-RNAi*]). ***p < 0.001, ****p < 0.0001. (**N**) Quantification of F-actin mean intensity in the genotypes shown in J-L. Statistical significance was determined using a Brown-Forsythe and Welch ANOVA test (n = 31 [*ap>LacZ*], n = 22 [*ap>Yki*], n = 19 [*ap>Yki, dPGC1-RNAi*]). *p < 0.05, **p < 0.01. **(O, P)** Confocal micrographs showing magnifications from tumorous wing discs with the following genotypes: *ap-Gal4, UAS-Yki, UAS-GFP, UAS-LacZ* (O) and *ap-Gal4, UAS-Yki, UAS-GFP, UAS-dPGC1-RNAi* (P). PH3 labels mitotic cells and is shown in red. GFP is shown in green. DAPI labels the DNA and is shown in blue. Scale bars, 10 μm. (**Q**) Quantification of number of PH3-positive cells per GFP-positive area in the genotypes shown in O and P. Statistical significance was determined using an unpaired t-test (n = 23 [*ap>Yki*], n = 19 *[ap>Yki, dPGC1-RNAi*]). ****p < 0.0001.

To complement the overexpression studies and assess the role of dPGC1 in a more physiologically relevant context, we analyzed *Warts* (*Wts*) mutant clones. Wts (Lats in mammals) is a core kinase in the Hippo pathway that phosphorylates and inhibits Yki, preventing its nuclear localization. As previously reported, loss of *Wts* mimics *Yki* overexpression and results in tissue overgrowth (Huang *et al*., 2005). Consistent with this, *Wts* mutant clones were larger than control clones (Fig 2F, G, I). Notably, *Wts* mutant clones expressing *dPGC1-RNAi* exhibited even greater overgrowth (Fig 2G-I), supporting the idea that dPGC1 acts to restrain tissue expansion in cells with elevated Yki activity. These results validate our previous observations and further support a role for dPGC1 in limiting Yki-driven tissue overgrowth.

To determine whether the enhanced growth phenotype observed upon *dPGC1* depletion could be explained by increased *Yki* expression or activity, we used quantitative PCR (qPCR) to measure the transcript levels of *Yki* and its target genes *Cyclin E*, *Diap1*, and *bantam* in wing imaginal discs with or without *dPGC1* knockdown. These analyses revealed no significant changes, suggesting that the phenotype is not driven by elevated *Yki* expression or transcriptional activity (Fig S3).

Given that dPGC1 modulates Yki-driven overgrowth, we asked whether it acts as a general regulator of oncogenic growth or plays a more specific role in restraining Yki-induced tumorigenesis. To address this, we downregulated *dPGC1* in tissues overexpressing the oncogenes *Epidermal Growth Factor Receptor* (*EGFR*) or the *Insulin Receptor* (*InR*). In both cases, *dPGC1* depletion did not enhance tissue overgrowth, in contrast to the effect observed in *Yki*-overexpressing discs (compare Fig S4 with Fig 2C-E). This indicates that the tumor-suppressive role of dPGC1 is specifically relevant in the context of Yki-driven tumorigenesis rather than being a general response to oncogenic signaling.

### dPGC1 Restrains Oncogenic Traits in Yki-Induced Tumorous Discs

Having established that dPGC1 limits Yki-driven tissue overgrowth, we next asked whether it also suppresses other oncogenic traits associated with tumorigenesis. In addition to increased tissue size, malignant fly tumors often express the secreted matrix metalloproteinase 1 (Mmp1), which promotes cell migration and invasion by degrading the basement membrane (Beaucher *et al*, 2007; Uhlirova & Bohmann, 2006). While we did not detect Mmp1 expression in control discs, Mmp1 was observed in discrete patches of cells in discs overexpressing *Yki* (Fig 2J, K, M). Notably, discs co-expressing *Yki* and *dPGC1-RNAi* displayed a robust increase in Mmp1 (Fig 2K-M). *Mmp1* is a target gene of the c-Jun N-terminal kinase (JNK) pathway, a signaling cascade activated by internal and external stressors that regulates proliferation, apoptosis, and cell migration (La Marca & Richardson, 2020; Pinal *et al*, 2019). JNK acts as an oncogenic partner of pro-tumorigenic factors such as Ras^V12^ and Yki (Igaki *et al*, 2006; Uhlirova & Bohmann, 2006). In these contexts, JNK induces the accumulation of F-actin, which is observed in cells undergoing malignant transformation (Enomoto *et al*, 2015; Fernandez *et al*, 2011; Fernandez *et al*, 2014; George & Jacobs, 2019; Khoo *et al*, 2013; Uhlirova & Bohmann, 2006). Consistent with these findings, we observed elevated F-actin levels in *Yki + dPGC1-RNAi* tumorous discs, along with increased expression of the JNK target gene *Mmp1* (Fig 2J-L, N). These results were confirmed using an independent UAS-driven *dPGC1-shRNA* line (Fig S5), whose knockdown efficiency is shown in Fig S2.

In addition to invasive and structural changes, uncontrolled proliferation is a hallmark of oncogenic growth (Hanahan & Weinberg, 2011). To determine whether this feature was also present in *Yki + dPGC1-RNAi* tumorous discs, we examined mitotic activity using phospho-Histone H3 (PH3) staining, a well-established marker of cells undergoing mitosis and thus a proxy for cell proliferation. This analysis revealed a significant increase in mitotic cells, indicating elevated proliferative activity and highlighting an additional oncogenic feature of these tumors (Fig 2O-Q). These findings suggest that *dPGC1* depletion not only enhances tissue overgrowth and promotes markers of malignancy but also accelerates cell cycle progression in the context of Yki activation.

In sum, our results show that *dPGC1* knockdown enhances Yki-induced overgrowth, leading to the formation of tumorous discs that exhibit markers of malignancy. Although *dPGC1* depletion does not have a major impact on normal growth, it is required to maintain controlled growth and tissue integrity in the context of *Yki* upregulation.

### Mitochondrial Membrane Potential in Yki-Driven Tumors

Given the central role of dPGC1 in maintaining mitochondrial activity, we sought to evaluate mitochondrial functionality in tumors driven by Yki activation and *dPGC1* depletion. As a surrogate for mitochondrial function, we measured mitochondrial membrane potential (MMP). MMP is a key indicator of mitochondrial function, reflecting the integrity of the electron transport chain and the capacity for ATP production via oxidative phosphorylation. To evaluate MMP, we employed tetramethylrhodamine ethyl ester (TMRE), a cell-permeant fluorescent dye that accumulates in mitochondria in proportion to their electrochemical gradient (Ehrenberg *et al*, 1988). TMRE staining showed no significant difference in MMP between discs expressing *Yki* alone and those co-expressing *Yki* with *dPGC1-RNAi* (Fig S6). These results indicate that *dPGC1* depletion does not impair mitochondrial membrane potential in Yki-driven tumors.

### dPGC1 Loss Selectively Upregulates Mitochondrial Fusion Genes in Yki-Driven Tumors

Given that mitochondrial membrane potential was not significantly affected by *dPGC1* depletion in Yki-driven tumors, we next investigated whether this genetic interaction alters the transcriptional landscape of genes involved in mitochondrial dynamics. This analysis was motivated by previous findings showing that PGC1 family proteins regulate the expression of genes involved in mitochondrial biogenesis and function (Villena, 2015) and that Yki modulates mitochondrial architecture to support tissue growth (Nagaraj *et al*., 2012). We analyzed whether *dPGC1* downregulation in the context of Yki activation affects the expression of genes involved in these processes. By using qPCR, we compared the mRNA levels of genes involved in different aspects of mitochondrial dynamics such as mitochondrial biogenesis, mitochondrial fusion and fission, mitophagy, and mitochondrial transport (Archer, 2013). Among genes controlling mitochondrial biogenesis, we quantified the expression of *estrogen-related receptor* (*dERR*), *Ets at 97D* (*Ets97D*), and *erect wing* (*ewg*) (Baltzer et al, 2009; Chen et al, 2008; Misra et al, *2017*; Rai et al, 2014; Takata et al, 2001). We also assessed the expression of the genes *dMfn* and *Opa1*, which control fusion between outer and inner membranes, respectively; and *Dynamic-related protein 1* (*Drp1*), which mediates mitochondrial fission (Deng et al, 2008). Additionally, we quantified the expression of genes involved in mitophagy, including *PTEN-induced putative kinase 1* (*Pink1*), *parkin* (*park*), and *Autophagy-related 7* (*Atg7*) (Narendra et al, 2010; Vincow et al, 2013). Finally, we measured the levels of expression of genes involved in mitochondrial transport, such as *milton* (*milt*) and *mitochondrial Rho* (*miro*) (Guo et al, 2005; Stowers et al, 2002). We analyzed four independent biological replicates where we compared mRNA levels between *Yki*-upregulating discs (control) and *Yki* + *dPGC1-RNAi* tumorous discs (experimental condition). We found that discs co-expressing *Yki* and *dPGC1-RNAi* showed a significant increase in the expression of *dMfn* and *Opa1*, GTPases promoting mitochondrial fusion (Fig 3A). Interestingly, *milt* and *miro* were also upregulated in *Yki + dPGC1-RNAi* tumorous discs (Fig 3A). Although miro and milt play central roles in mitochondrial axonal transport, previous reports have suggested that they can also affect mitochondrial morphology (Ding *et al*, 2016; Liu & Hajnoczky, 2009; Panchal & Tiwari, 2020; Tang, 2015). For instance, *miro* overexpression in *Drosophila* models of Alzheimer’s disease has been reported to cause an increased mitochondrial average length (Panchal & Tiwari, 2020).

**Figure 3.**
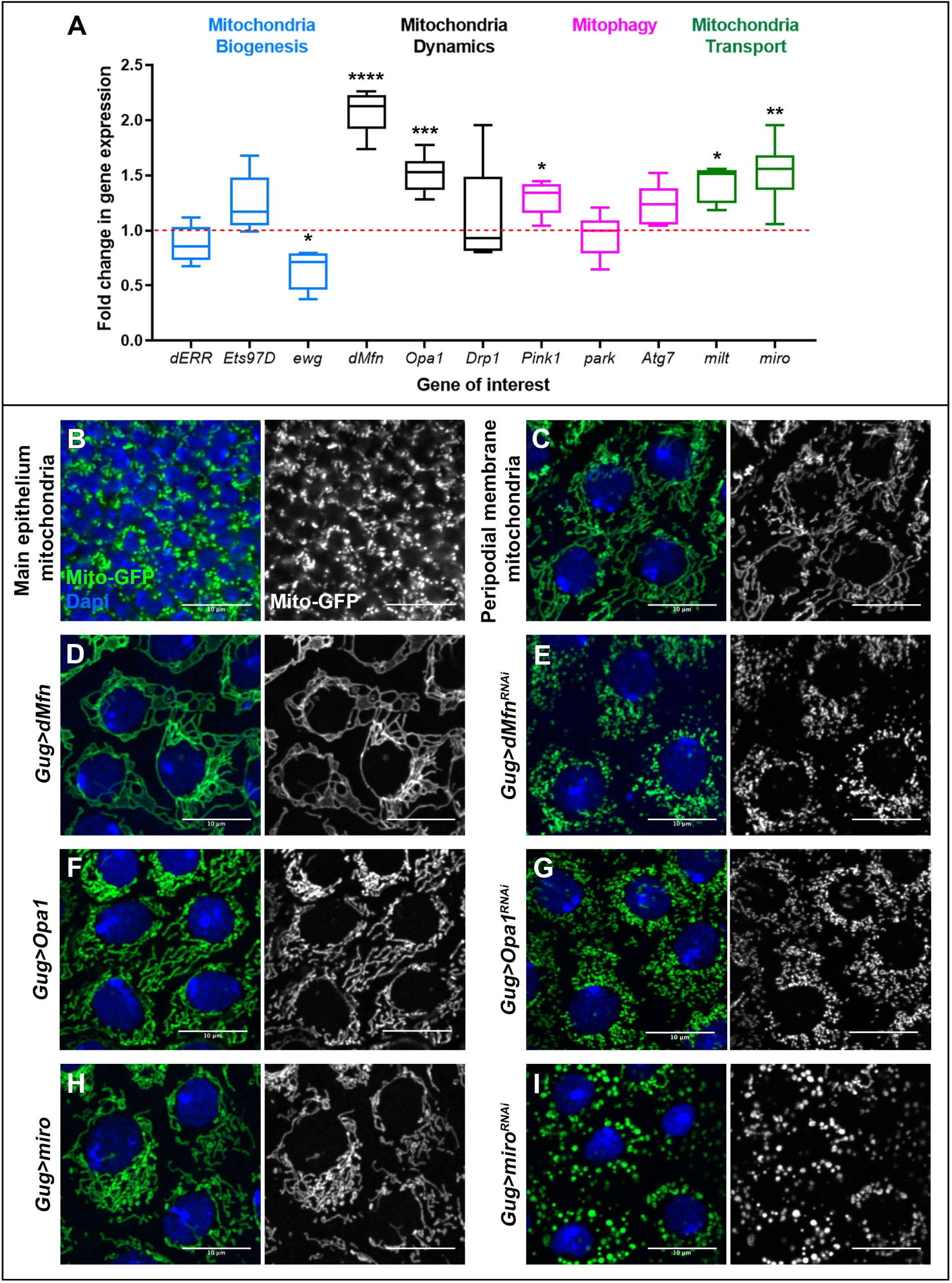
Aberrant Mitochondrial Dynamics in *Yki + dPGC1-RNAi* Tumorous Discs. **(A)** mRNA quantification by qPCR of the indicated genes in *ap-Gal4, UAS-GFP, UAS-Yki, UAS-dPGC1-RNAi* wing imaginal discs using *ap-Gal4, UAS-GFP, UAS-Yki, UAS-LacZ* as the control genotype. Genes are separated in different categories: mitochondria biogenesis (blue), mitochondria dynamics (black), mitophagy (pink), and mitochondria transport (green). Each genotype was run in five to six biological replicates. RP49 was used as the housekeeping gene. Statistical significance was determined using unpaired t-tests with Welch’s correction for parametric data and Mann-Whitney tests for non-parametric data. *p < 0.05, **p < 0.01, ***p < 0.001, ****p < 0.0001. **(B)** Confocal micrograph of the main epithelium of an *ap-Gal4, UAS-Mito-GFP* wing imaginal disc. Mitochondria are labeled with Mito-GFP and are shown in green. DAPI is shown in blue. Scale bar, 10 μm. **(C)** Confocal micrograph of the peripodial membrane of a *Gug-Gal4, UAS-Mito-GFP* wing imaginal disc. Mitochondria are labeled with Mito-GFP and are shown in green. DAPI is shown in blue. Scale bar, 10 μm. **(D-I)** Confocal micrographs of the peripodial membrane in the following genotypes: *Gug-Gal4, UAS-Mito-GFP, UAS-dMfn* (D); *Gug-Gal4, UAS-Mito-GFP, UAS-dMfn-RNAi* (E); *Gug-Gal4, UAS-Mito-GFP, EP-Opa1* (F); *Gug-Gal4, UAS-Mito-GFP, UAS-Opa1-RNAi* (G); *Gug-Gal4, UAS-Mito-GFP, UAS-miro* (H); and *Gug-Gal4, UAS-Mito-GFP, UAS-miro-RNAi* (I). Mitochondria are labeled with Mito-GFP and are shown in green. DAPI is shown in blue. Scale bars, 10 μm.

*dPGC1* depletion in otherwise normal discs did not affect the expression of genes controlling fusion/fission mechanisms to the same extent as observed in a context of *Yki* upregulation (Fig S7). This observation suggests that the transcriptional response of mitochondrial dynamics genes to *dPGC1* depletion is influenced by Yki activity, indicating that mitochondrial remodeling becomes particularly responsive to *dPGC1* loss in an oncogenic environment driven by *Yki* upregulation.

To further explore the relationship between dPGC1 and mitochondrial fusion in the context of *Yki* upregulation, we analyzed the expression of *dMfn* and *Opa1* in discs overexpressing *dPGC1* alongside *Yki*. In contrast to the effects observed upon *dPGC1* downregulation, *dPGC1* upregulation in *Yki*-overexpressing discs did not significantly affect tissue growth and did not alter *dMfn* and *Opa1* mRNA levels (Fig S8). These findings suggest that the transcriptional upregulation of mitochondrial fusion genes is a specific consequence of *dPGC1* knockdown in a Yki-activated context, highlighting a sensitized mitochondrial response to Yki signaling rather than a general effect of altered dPGC1 levels.

### Functional Analysis of Mitochondrial Fusion Genes in Wing Disc Development

The genes *dMfn, Opa1, milt,* and *miro*, which are upregulated in *Yki + dPGC1-RNAi* tumorous discs, have been extensively studied in various model systems (Archer, 2013; Chan, 2020; Lee & Lu, 2014; Zorzano *et al*, 2010). However, their specific functions in regulating mitochondrial morphology in the *Drosophila* wing imaginal disc remain uncharacterized. To address this, we experimentally manipulated the expression of these genes during wing disc development. To visualize mitochondria, we used a mitochondrial-targeted GFP (Mito-GFP) transgene, which contains localization sequences that direct GFP to the mitochondria (Rizzuto *et al*, 1995).

The wing imaginal disc is composed of two distinct epithelial layers: a densely packed columnar epithelium, referred to as the main epithelium, and a squamous epithelium known as the peripodial membrane (Gibson & Schubiger, 2001). The compact architecture and limited cytoplasmic volume of the main epithelium pose challenges for high-resolution imaging of subcellular structures such as mitochondria (Fig 3B). To circumvent these limitations, we performed mitochondrial imaging in the peripodial membrane, which consists of large, flattened cells with expanded cytoplasmic space that facilitates clear visualization of mitochondrial morphology. In this context, mitochondria appeared as distinct and well-defined organelles (Fig 3C), enabling a reliable assessment of morphological changes following genetic manipulations.

To assess the roles of genes upregulated in *Yki + dPGC1-RNAi* tumors, we used the Grunge-Gal4 (Gug-Gal4) driver to manipulate *dMfn, Opa1*, and *miro* specifically in the peripodial membrane of the wing imaginal disc (Gibson *et al*, 2002). This allowed us to visualize mitochondrial changes resulting from their experimental deregulation. dMfn and Opa1 are central regulators of mitochondrial fusion. Upregulation of *dMfn* led to a connected mitochondrial network, indicative of enhanced fusion (Fig 3D; quantified in Fig S9). In contrast, overexpression of *Opa1* produced a subtler phenotype and mitochondria appeared slightly elongated compared to controls, but the overall impact on network connectivity was modest relative to *dMfn* overexpression (compare Fig 3F with Fig 3D; quantified in Fig S9). Depletion of *dMfn1* or *Opa1* led to mitochondrial fragmentation (Fig 3E, G; quantified in Fig S9). The phenotypic differences observed when *dMfn* and *Opa1* were overexpressed likely reflect the distinct molecular roles of dMfn and Opa1 within the mitochondrial fusion machinery. While both proteins promote fusion, dMfn primarily facilitates the merging of the outer mitochondrial membrane, a process that directly contributes to the formation of interconnected networks. Opa1 regulates inner membrane fusion and cristae architecture, which may lead to elongation without extensive network formation.

*milt* and *miro* were also upregulated in *Yki + dPGC1-RNAi* tumors. Although milt and miro are traditionally associated with mitochondrial transport, recent studies suggest their involvement in mitochondrial dynamics (Ding *et al*., 2016; Liu & Hajnoczky, 2009; Panchal & Tiwari, 2020; Tang, 2015). In our expression analysis, *miro* showed the highest level of upregulation between the two (Fig 3A), making it a particularly compelling candidate for functional investigation. Altering *miro* expression led to noticeable changes in mitochondrial morphology (Fig 3H, I; quantified in Fig S9). *miro* overexpression resulted in mitochondria that appeared elongated but not as extensively fused as those observed in *dMfn*-overexpressing imaginal discs (Fig 3H; quantified in Fig S9). The most pronounced effects were observed upon *miro* depletion. In these cells, mitochondria appeared smaller and more condensed, as indicated by the increased intensity and compaction of the Mito-GFP signal (Fig 3I; quantified in Fig S9). This phenotype differs qualitatively from the fragmentation seen with *dMfn* or *Opa1* knockdown and may reflect a different role in mitochondrial morphology. Given its established function in mitochondrial transport, *miro* downregulation may disrupt mitochondrial positioning and cytoskeletal interactions, leading to clustering rather than classical fragmentation.

The efficiency of the *UAS-RNAis* used to knock down *dMfn, Opa1*, and *miro* is shown in Fig S2.

### Enhanced Mitochondrial Fusion in Yki-Driven Tumors Upon *dPGC1* Depletion

Our analyses have revealed transcriptional upregulation of mitochondrial fusion genes in *Yki + dPGC1-RNAi* tumors, suggesting a shift in mitochondrial dynamics. To investigate whether these transcriptional changes translate into altered mitochondrial morphology, we examined mitochondria in the tumorous tissue. Since the ap-Gal4 driver used to induce tumor formation is active in the main epithelium, which contains small, densely packed cells with limited cytoplasmic space, confocal imaging of mitochondria in this tissue proved technically challenging (Fig 3B). To overcome this limitation and obtain high-resolution structural information, we employed electron microscopy (EM) to assess mitochondrial morphology in Yki-driven tumors with or without *dPGC1* depletion.

Consistent with the upregulation of mitochondrial fusion genes such as *dMfn* and *Opa1*, we observed that mitochondria in *Yki + dPGC1-RNAi* tumorous discs were larger and more elongated than those in *Yki*-expressing discs (Fig 4A, B, D, E). Next, we assessed the consequences of downregulating *dMfn*, the mitochondrial fusion gene showing the highest upregulation in *Yki + dPGC1-RNAi* tumorous discs (Fig 3A). We found that mitochondria in tumors with reduced *dMfn* appeared larger and more rounded compared to those in *Yki + dPGC1-RNAi* tumors (Fig 4B-E). Notably, these swollen mitochondria exhibited disrupted cristae, with some displaying a characteristic onion-like ultrastructure (Fig 4C, C’). In addition, we observed vesicle-like structures in those mitochondria, further suggesting that cristae organization and mitochondrial integrity were compromised upon *dMfn* depletion (Fig 4C, C’). These structural abnormalities suggest that cristae disorganization may impair mitochondrial function, potentially disrupting energy production and cellular homeostasis, which could in turn limit the ability of cells to sustain oncogenic growth.

**Figure 4.**
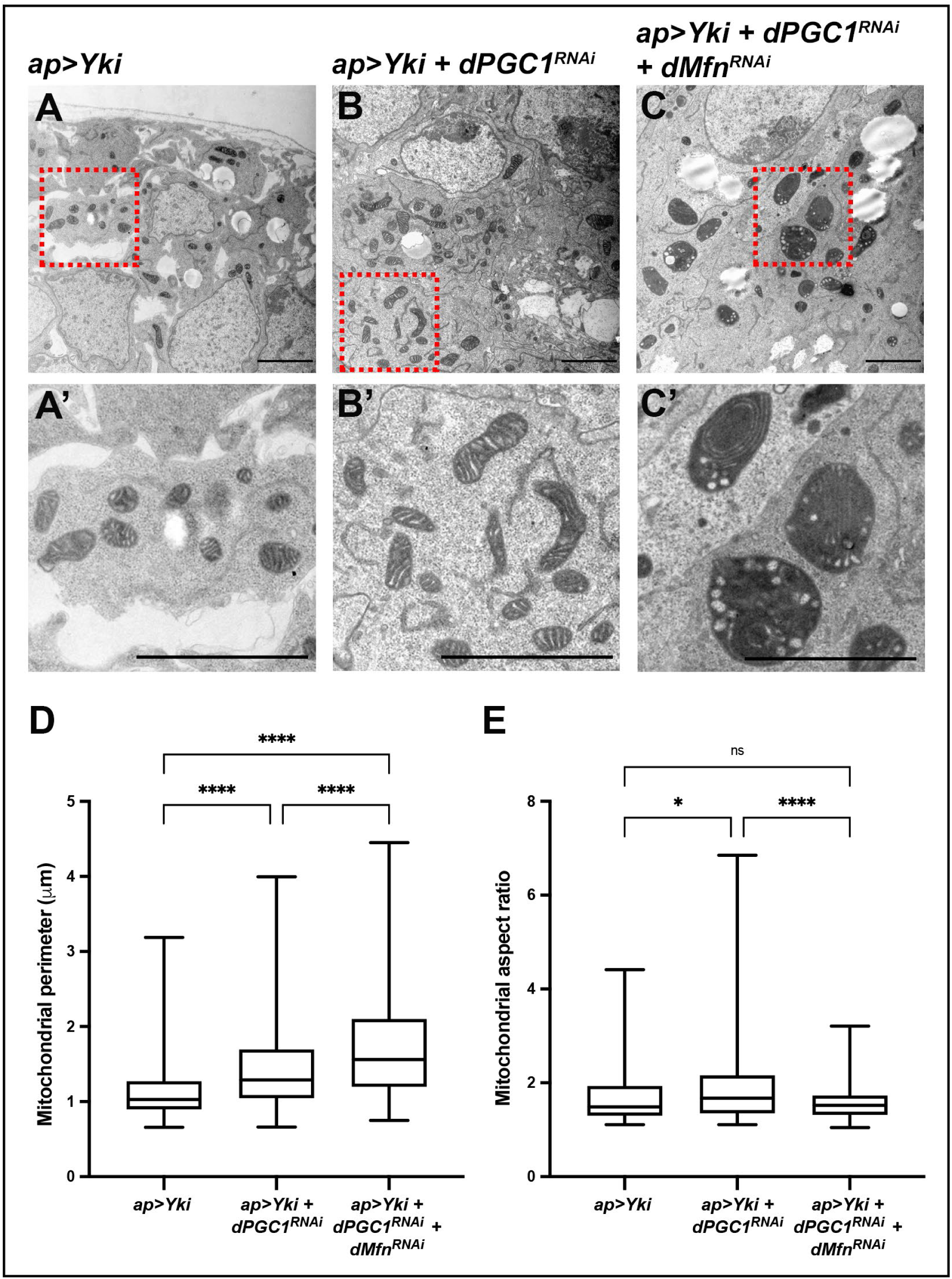
dPGC1 and dMfn Regulate Mitochondrial Morphology in Yki-Driven Tumors. **(A-C)** Representative electron microscopy micrographs obtained from discs of the following genotypes: *ap-Gal4, UAS-GFP, UAS-Yki, UAS-LacZ* (A); *ap-Gal4, UAS-GFP, UAS-Yki, UAS-dPGC1-RNAi* (B); and *ap-Gal4, UAS-GFP, UAS-Yki, UAS-dPGC1-RNAi, UAS-dMfn-RNAi* (C). The dashed red squares in A-C indicate the regions that are shown as magnifications in A’-C’. Scale bars, 2 μm. **(D, E)** Quantification of mitochondrial perimeter (D) and mitochondrial aspect ratio (E) in the genotypes shown in A-C. Statistical significance was determined using one-way ANOVA tests (n = 164 [*ap>Yki*], n = 214 [*ap>Yki, dPGC1-RNAi*], n = 218 [*ap>Yki, dPGC1-RNAi, dMfn-RNAi*]). ns, non-significant. *p < 0.05, ****p < 0.0001.

### Contribution of Mitochondrial Fusion Genes to Yki-Mediated Tumor Growth

Previous studies have shown that Yki upregulates genes involved in mitochondrial fusion, a process that plays a central role in its growth-promoting activity (Nagaraj *et al*., 2012). Building on this, we demonstrate that depletion of *dPGC1* further enhances Yki-induced tissue overgrowth, with the resulting tumorous discs exhibiting elongated mitochondria. These findings led us to hypothesize that the upregulation of mitochondrial fusion genes may contribute to tumor formation when *Yki* is overexpressed and *dPGC1* is knocked down. If this is the case, then reducing the expression of these fusion-promoting genes should mitigate tumor growth. To test this, we used *UAS-RNAi* transgenes targeting *dMfn* or *Opa1* in *Yki + dPGC1*-depleted tumors, which significantly reduced tissue growth compared to control tumors (Fig 5A–D, G). These results indicate that mitochondrial fusion supports tumor expansion in this context. Next, we investigated whether upregulation of mitochondrial fusion genes alone is sufficient to drive tumorigenesis in cooperation with the proto-oncogene Yki. Co-expression of *Yki* with either *dMfn* or *Opa1* led to a marked increase in disc size (Fig 5A, E–G), indicating that enhanced mitochondrial fusion can potentiate Yki-driven tissue overgrowth. Importantly, these tumors exhibited hallmark features of malignancy, including elevated expression of *Mmp1* and disrupted epithelial architecture, as revealed by F-actin staining (Fig 5H, I). In contrast, knockdown or overexpression of these fusion genes in otherwise normal discs resulted in only minor changes in tissue size, effects that were significantly less pronounced than those observed in the tumor context (Fig S10). Our findings highlight that dMfn and Opa1 are critical modulators of Yki-driven tumorigenesis.

**Figure 5.**
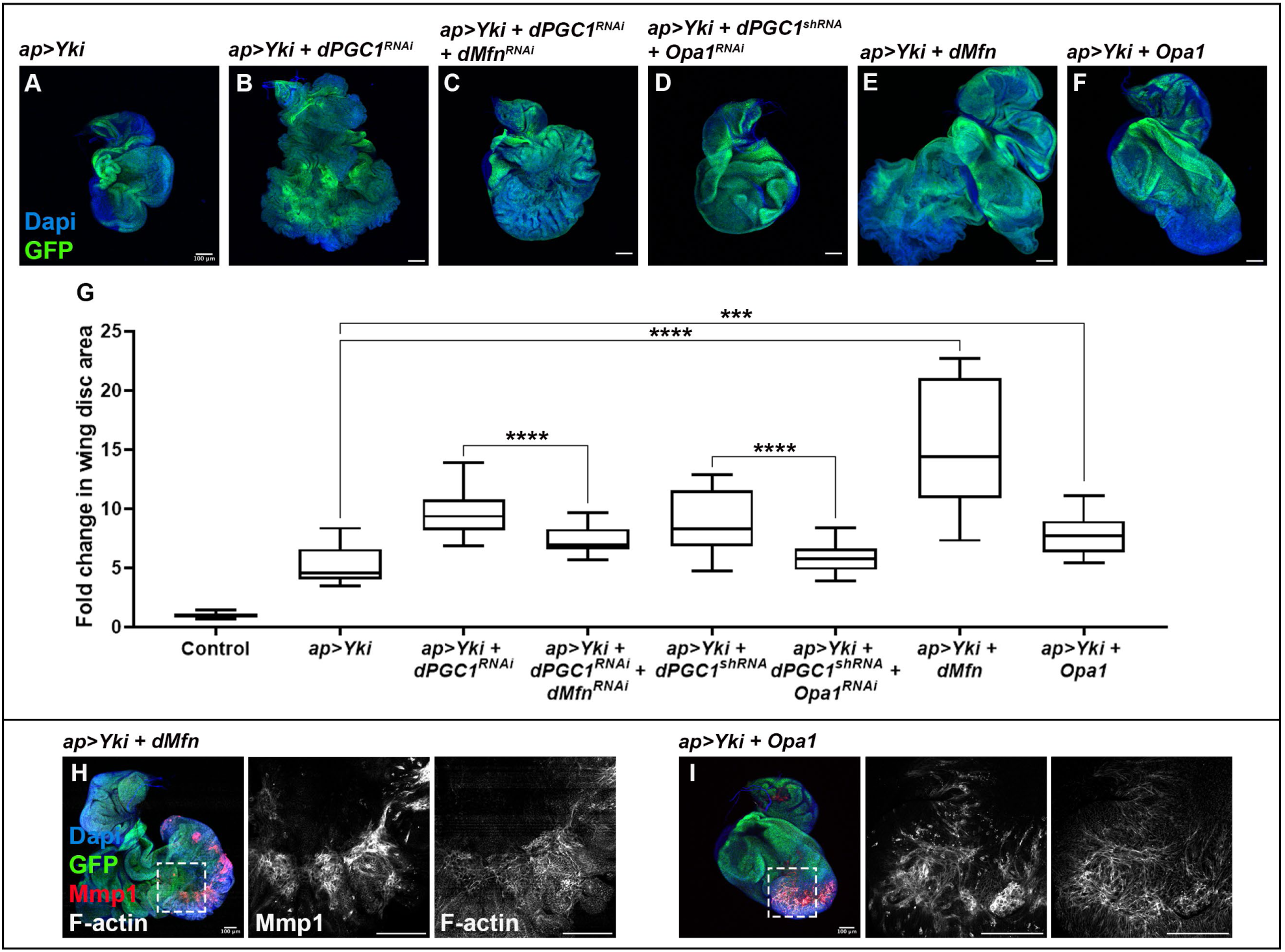
dMfn and Opa1 Synergize with Yki to Promote Tumor Growth and Malignancy. **(A-F)** Confocal micrographs showing discs of the following genotypes: *ap-Gal4, UAS-GFP, UAS-Yki, UAS-LacZ* (A); *ap-Gal4, UAS-GFP, UAS-Yki, UAS-dPGC1-RNAi* (B); *ap-Gal4, UAS-GFP, UAS-Yki, UAS-dPGC1-RNAi, UAS-dMfn-RNAi* (C); *ap-Gal4, UAS-GFP, UAS-Yki, UAS-dPGC1-shRNA, UAS-Opa1-RNAi* (D); *ap-Gal4, UAS-GFP, UAS-Yki, UAS-dMfn* (E); and *ap-Gal4, UAS-GFP, UAS-Yki, EP-Opa1* (F). GFP is shown in green, and DAPI labels the DNA and is shown in blue. Scale bars, 100 μm. **(G)** Quantification of GFP-positive area in third instar wing imaginal discs of the genotypes shown in A-F. GFP-positive areas are normalized to the mean of the control (*ap>LacZ*). Statistical significance was determined using unpaired t-tests with Welch’s correction (n = 18 [*ap>LacZ*], n = 20 [*ap>Yki*], n = 20 [*ap>Yki, dPGC1-RNAi*], n = 20 [*ap>Yki, dPGC1-RNAi, dMfn-RNAi*], n = 20 [*ap>Yki, dPGC1-shRNA*], n = 20 [*ap>Yki, dPGC1-shRNA, Opa1-RNAi*], n = 13 [*ap>Yki, dMfn*], n = 12 [*ap>Yki, Opa1*]). ***p < 0.001, ****p < 0.0001. **(H, I)** Confocal micrographs of imaginal discs of the following genotypes: *ap-Gal4, UAS-GFP, UAS-Yki, UAS-dMfn* (H) and *ap-Gal4, UAS-GFP, UAS-Yki, UAS-Opa1* (I). F-actin labels cell polarity and epithelial organization and is shown in grayscale. Mmp1 is shown in red. GFP is shown in green. DAPI labels the DNA and is shown in blue. Scale bars, 100 μm.

To further explore the oncogenic contribution of the genes upregulated in our expression analysis (Fig 3A), we examined the function of miro. We found that, in contrast to *dMfn* and *Opa1*, *miro* upregulation was not sufficient to drive tumorigenesis in combination with Yki. Moreover, *miro* depletion did not alter the growth of *Yki + dPGC1-RNAi* tumors (Fig S11).

In summary, our results demonstrate that genes controlling mitochondrial morphology play a gene-specific and context-dependent role in Yki-mediated tumorigenesis. While upregulation of fusion genes such as *dMfn* and *Opa1* synergizes with *Yki* to promote tissue overgrowth and malignancy, *miro* does not exhibit the same oncogenic potential.

### Upregulation of Cyclin E in Tumors Caused by *Yki* upregulation and *dPGC1* Depletion

Mounting evidence has established a connection between mitochondrial dynamics and the cell cycle (Ma *et al*, 2020). Proper control of the cell cycle is essential for accurate genome transmission during cell division, and its disruption can lead to genomic instability, which is a hallmark of cancer. (Hanahan, 2022; Hanahan & Weinberg, 2011). Cyclin-dependent kinases (CDKs) and their regulatory cyclins are well-established drivers of cell cycle progression (Besson *et al*, 2008; Satyanarayana & Kaldis, 2009). Beyond these canonical regulators, mitochondria have emerged as key modulators of cell cycle control, with their dynamic behavior contributing to the maintenance of genomic stability (Horbay & Bilyy, 2016; Lopez-Mejia & Fajas, 2015; Mandal *et al*, 2005; Owusu-Ansah *et al*, 2008).

Studies in mammalian cells and in *Drosophila* have shown that cells with highly fused mitochondria present an elevation in the levels of the oncogene *Cyclin E* (Finkel & Hwang, 2009; Mitra, 2013; Mitra *et al*, 2012; Mitra *et al*, 2009; Qian *et al*., 2012). Cyclin E peaks at the G1-S transition, activating CDK2 and promoting S phase entry (Siu *et al*, 2012). Aberrant Cyclin E levels can trigger premature entry in S phase, leading to replicative stress and DNA damage, which are key contributors to genomic instability in cancer (Gaillard *et al*, 2015; Molano-Fernandez *et al*, 2022; Saxena & Zou, 2022).

We have shown that *dPGC1* depletion in epithelial cells with elevated Yki signaling leads to an increase in mitochondrial size, which correlates with the transcriptional upregulation of genes promoting mitochondrial fusion, and is associated with tumor formation. These observations prompted us to investigate whether *Yki + dPGC1-RNAi* tumors exhibit abnormal levels of Cyclin E. We first examined *cyclin E* mRNA levels and found no significant differences between discs expressing *Yki* alone and those co-expressing *Yki* with *dPGC1-RNAi* (Fig 6A). Interestingly, although transcript levels remained unchanged, Cyclin E protein levels were elevated in tumors driven by *Yki* overexpression and *dPGC1* depletion (Fig 6B, C; Fig S12). These results indicate that Cyclin E upregulation occurs post-transcriptionally and are consistent with previous studies linking mitochondria to Cyclin E regulation through non-transcriptional mechanisms (Mandal *et al*., 2005; Owusu-Ansah *et al*., 2008).

**Figure 6.**
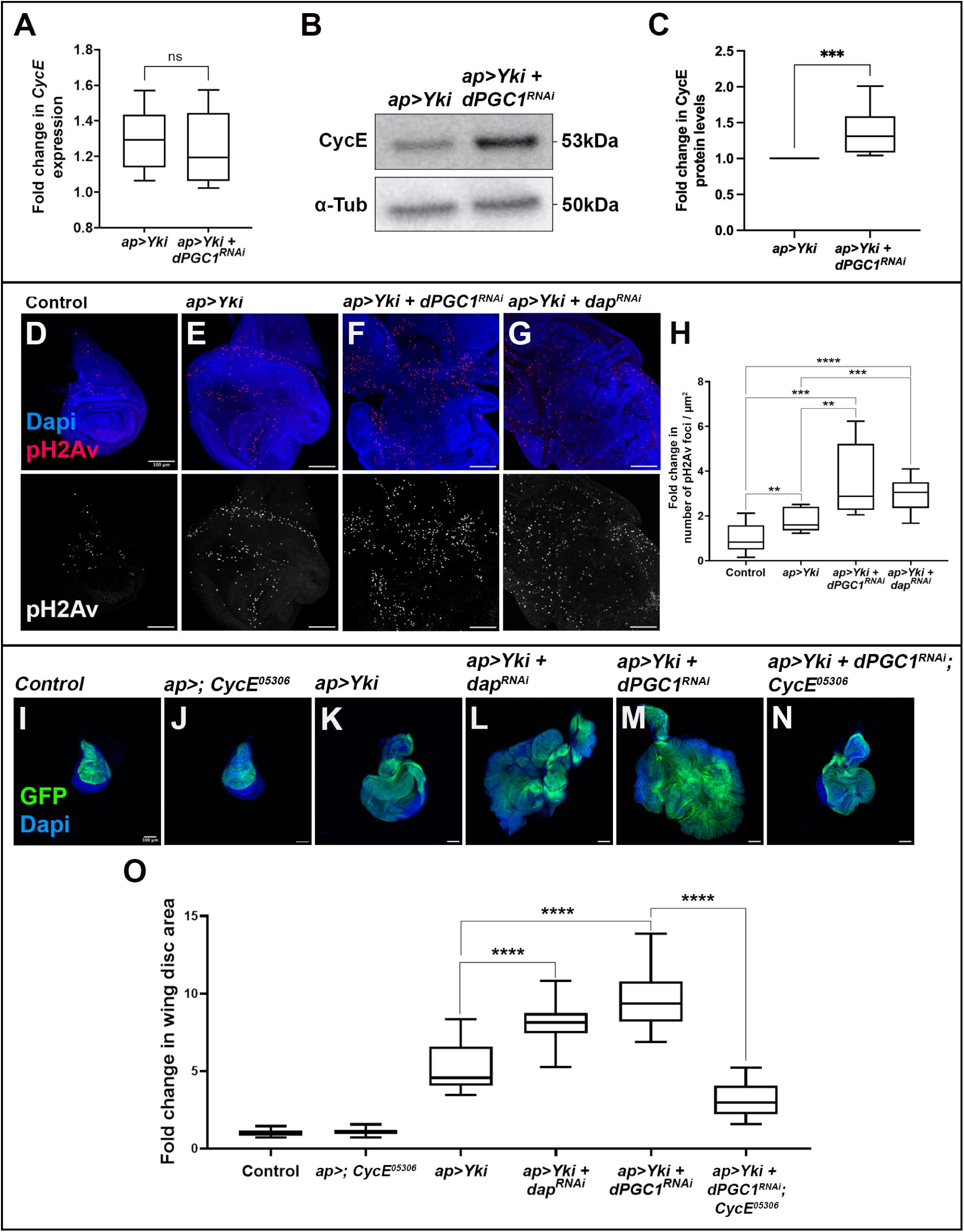
Cyclin E Stabilization Drives Tumor Growth and DNA Damage in *Yki + dPGC1-RNAi* Discs. **(A)** Quantification of *Cyclin E* mRNA by qPCR in imaginal discs of the following genotypes: *ap-Gal4, UAS-GFP, UAS-Yki, UAS-LacZ* (*ap>Yki*) and *ap-Gal4, UAS-GFP, UAS-Yki, UAS-dPGC1-RNAi* (*ap>Yki, dPGC1-RNAi*). Each genotype was run in four to five biological replicates. RP49 was used as the housekeeping gene. Statistical significance was determined using an unpaired t-test with Welch’s correction. ns, non-significant. **(B)** Western blot showing Cyclin E protein levels in third instar wing imaginal discs of the indicated genotypes. α-Tubulin was used as the loading control. (**C**) Relative quantification of Cyclin E protein levels in wing imaginal discs of the genotypes shown in B. Cyclin E protein levels were normalized to α-Tubulin protein levels for each western blot membrane. The obtained Cyclin E/α-Tubulin ratio was normalized to that of the control (*ap>Yki*). Statistical significance was determined using a Mann-Whitney test (n = 7). ***p < 0.001. **(D-G)** Maximum projection of confocal micrographs of imaginal discs of the following genotypes: *ap-Gal4, UAS-GFP, UAS-LacZ* (D); *ap-Gal4, UAS-GFP, UAS-Yki, UAS-LacZ* (E); *ap-Gal4, UAS-GFP, UAS-Yki, UAS-dPGC1-RNAi* (F); and *ap-Gal4, UAS-GFP, UAS-Yki, UAS-dap-RNAi* (G). pH2Av is shown in red, and DAPI is shown in blue. Scale bars, 100 μm. **(H)** Quantification of pH2Av-positive foci in the genotypes shown in D-G. Data was normalized to the mean of the control (*ap>LacZ*). Statistical significance was determined using unpaired t-tests with Welch’s correction (n = 12 [*ap>LacZ*], n = 12 [*ap>Yki*], n = 12 [*ap>Yki, dPGC1-RNAi*], n = 20 [*ap>Yki, dap-RNAi*]). **p < 0.01, **p < 0.001, ****p < 0.0001. **(I-N)** Confocal micrographs showing discs of the following genotypes: *ap-Gal4, UAS-GFP, UAS-LacZ* (I); *ap-Gal4, UAS-GFP/Cyclin E-05306* (J); *ap-Gal4, UAS-GFP, UAS-Yki, UAS-LacZ* (K); *ap-Gal4, UAS-GFP, UAS-Yki, UAS-dap-RNAi* (L), *ap-Gal4, UAS-GFP, UAS-Yki, UAS-dPGC1-RNAi* (M); and *ap-Gal4, UAS-GFP, UAS-Yki, UAS-dPGC1-RNAi/Cyclin E-05306* (N). GFP is shown in green, and DAPI labels the DNA and is shown in blue. Scale bars, 100 μm. **(O)** Quantification of GFP-positive area in third instar wing imaginal discs of the genotypes shown in I-N. GFP-positive area is normalized to the mean of the control (*ap>LacZ*). Statistical significance was determined using unpaired t-tests with Welch’s correction (n = 18 [*ap>LacZ*], n = 20 [*ap>; CycE-05306*], n = 20 [*ap>Yki*], n = 20 [*ap>Yki, dap-RNAi*] n = 20 [*ap>Yki, dPGC1-RNAi*], n = 20 [*ap>Yki, dPGC1-RNAi; CycE-05306*]). ****p < 0.0001.

Aberrant Cyclin E levels can lead to replicative stress and DNA damage in human cells and *Drosophila* (Ekholm-Reed *et al*, 2004; Molano-Fernandez *et al*., 2022; Spruck *et al*, 1999). In response to DNA damage, checkpoint proteins phosphorylate the histone H2AX (H2Av in *Drosophila*) to mark the sites of damage (Blackford & Jackson, 2017; Fernandez-Capetillo *et al*, 2003). Antibodies recognizing the phosphorylated H2Av (pH2Av) can serve as markers to detect DNA damage. As expected, *Yki* overexpression led to an increase in pH2Av-positive foci (Fig 6D, E, H). Notably, this effect was enhanced when *dPGC1* was knocked down (Fic 6E, F, H), which is consistent with an increase in Cyclin E. These observations revealed that *dPGC1* depletion in discs activating the *Yki* proto-oncogene leads to an increase in Cyclin E levels and DNA damage.

Oncogene-induced DNA damage is a mutagenic source in cancer, and DNA damage can promote tumorigenesis in *Drosophila* (Dekanty *et al*, 2015; Gerlach & Herranz, 2020; Halazonetis *et al*, 2008). We then analyzed whether Cyclin E upregulation contributes to the growth of *Yki + dPGC1-RNAi* tumors. To investigate the role of Cyclin E dosage in tumor growth, we induced the formation *Yki + dPGC1-RNAi* tumors in a *Cyclin E* heterozygous (*+/-*) background and analyzed whether reduced Cyclin E levels affected tumor development. We observed that *Cyclin E* heterozygous discs did not display any growth defects (Fig 6I, J, O). However, when *Yki* was overexpressed and *dPGC1* was depleted in a *Cyclin E* heterozygous background, tumor size was dramatically reduced, indicating that reduced *Cyclin E* dosage specifically impairs tumor growth under these conditions (Fig 6M, N, O). This suggests that Cyclin E upregulation is a central factor driving tumorigenesis in *Yki + dPGC1-RNAi* tumors.

Next, we asked whether upregulation of Cyclin E was sufficient to enhance Yki-driven growth. Cyclin E activity is negatively regulated by the Cyclin E-dependent kinase inhibitor p21/dacapo (dap), which represses Cyclin E/CDK2 function. Therefore, reduced levels of dap increase Cyclin E/CDK2 activity (de Nooij *et al*, 1996; Lane *et al*, 1996). To test this, we depleted *dap* in *Yki*-expressing discs and found that this was sufficient to significantly enhance tumor growth (Fig 6K, L, O), supporting the idea that elevated Cyclin E activity promotes Yki-driven tumorigenesis. Interestingly, similar to what we observed in the *Yki + dPGC1-RNAi* context, *Yki + dap-RNAi* tumors also exhibited increased levels of DNA damage (Fig 6E, G, H), suggesting that Cyclin E upregulation may contribute to genomic instability in these tumors. The efficiency of *UAS-dap-RNAi* is shown in Fig S13. In sum, these results reveal that *dPGC1* downregulation in the context of Yki activation leads to Cyclin E upregulation, which promotes the accumulation of DNA damage and drives increased tumor growth.

Finaly, to determine whether the increase in Cyclin E is a general consequence of *dPGC1* depletion or specific to the oncogenic context of Yki activation, we analyzed Cyclin E levels in *dPGC1^1^* homozygous mutant wing discs (-/-) in the absence of *Yki* overexpression. We did not observe any detectable changes in Cyclin E protein levels under these conditions (Fig S14). This indicates that *dPGC1* depletion alone is not sufficient to stabilize Cyclin E and further supports the notion that the oncogenic traits observed, such as Cyclin E accumulation, are specific to the context of *Yki* upregulation.

## Discussion

Our findings demonstrate that dPGC1 functions as a tumor suppressor by limiting Yki-driven tissue overgrowth in the *Drosophila* wing imaginal disc. Under normal conditions, *dPGC1* mutants exhibit only a minor growth defect, and altering dPGC1 levels during physiological wing development does not lead to overt phenotypes. This indicates that dPGC1 is largely dispensable for normal tissue growth but becomes critical under oncogenic stress, particularly when Yki is hyperactivated. These observations align with recent work by the Halder group, which shows that Yki/YAP transcriptional input is not required for normal growth but instead drives ectopic tissue expansion through the induction of an aberrant genetic program (Kowalczyk *et al*, 2022). Such aberrant activation represents a pathological stress condition, further supporting the idea that dPGC1 is required to maintain tissue integrity under oncogenic challenges.

This selective requirement becomes even more striking when considering its specificity. Although dPGC1 does not modulate tumor-like growth driven by other oncogenic signals such as EGFR or InR, it plays a crucial role in suppressing Yki-induced overgrowth. Importantly, this suppression occurs without altering Yki expression or activity, suggesting that dPGC1 acts downstream or in parallel to Yki signaling. In this context, dPGC1 influences the transcription of mitochondrial fusion genes, pointing to a role in modulating mitochondrial dynamics in response to Yki-driven oncogenic stress. This is consistent with previous findings that link changes in mitochondrial morphology to the growth-promoting effects of Yki (Nagaraj *et al*., 2012), reinforcing a model in which dPGC1 safeguards tissue integrity by regulating mitochondrial function under pathological conditions.

This context-dependent role is consistent with observations in mammalian systems. For example, PGC1α knockout mice develop normally, indicating that PGC1 family members are not required for normal development (Leone *et al*., 2005), However, under stress conditions, PGC1α mutant mice exhibit a range of physiological defects, highlighting the importance of PGC1α in maintaining cellular homeostasis during pathological challenges (Arany *et al*, 2005; Lin *et al*., 2004). A similar context-specific function has been described for PGC1α in oncogenic processes. Its expression is frequently downregulated in various tumor types, suggesting a tumor-suppressive role (Feilchenfeldt *et al*, 2004; Torrano *et al*., 2016; Watkins *et al*, 2004; Zhang *et al*, 2007). Conversely, other studies have shown that PGC1α can promote tumorigenesis and metastasis in certain cellular and tissue contexts (Bhalla *et al*., 2011; LeBleu *et al*, 2014). Together, these findings highlight the importance of the cellular context in determining the pro- or antitumorigenic roles of these transcriptional regulators.

Yki promotes tissue overgrowth by regulating gene expression through its interaction with the TEAD transcription factor Scalloped, inducing targets such as *Cyclin E* and mitochondrial fusion genes like *dMfn* and *Opa1* (Bandura & Edgar, 2008; Nagaraj *et al*., 2012; Shu & Deng, 2017; Tapon *et al*, 2002). Our data show that *dPGC1* depletion in the context of Yki hyperactivation leads to upregulation of mitochondrial fusion genes, enlarged mitochondria, and enhanced tumor growth. These observations align with previous studies showing that increased mitochondrial fusion supports proliferation in Ras-transformed fibroblasts and *Drosophila* stem cells (Dubal *et al*, 2022; Garcez *et al*, 2020; Yao *et al*, 2019). Importantly, this regulation appears to be context-dependent, as *dPGC1* depletion in otherwise normal discs has little or no detectable impact on the expression of mitochondrial fusion and fission genes. These findings suggest that dPGC1 dampens the ability of Yki to induce the expression of genes inducing mitochondrial fusion, thereby limiting oncogenic proliferation. Supporting this, reducing dMfn levels in *Yki + dPGC1-RNAi* discs disrupt mitochondrial morphology and impairs tumor growth. EM analyses of these discs reveal swollen mitochondria with disorganized cristae, structures essential for respiration and ATP production. Such defects likely compromise mitochondrial function and contribute to reduced tumor expansion. Beyond their roles in fusion, Opa1 and Mfn2 also regulate cristae architecture and mitochondrial-ER contacts, which are critical for apoptosis, oxidative phosphorylation, and calcium signaling (Frezza *et al*, 2006; Naon *et al*, 2023). While our results support a role for enhanced mitochondrial fusion in Yki-driven tumorigenesis, we acknowledge that mitochondrial enlargement may also result from reduced fission activity or imbalances in mitochondrial biogenesis and mitophagy. Future studies employing live-cell imaging and dynamic assays will be essential to fully characterize the mitochondrial remodeling events underlying oncogenic transformation in this context. These broader functions may help explain how altered mitochondrial structure and inter-organelle communication contribute to the oncogenic phenotype.

Alterations in mitochondrial structure and function can influence key cell cycle regulators, linking metabolic and structural changes to proliferative outcomes (Lopez-Mejia & Fajas, 2015; Mandal *et al*., 2005; Mitra, 2013; Mitra *et al*., 2009; Owusu-Ansah *et al*., 2008). Mitochondrial fusion has been shown to stabilize and increase Cyclin E levels, promoting G1–S progression (Finkel & Hwang, 2009; Mitra, 2013). Consistent with this, our results show that *dPGC1* depletion in the context of *Yki* upregulation leads to increased Cyclin E protein levels. This correlates with elevated DNA damage, a hallmark of genomic instability commonly observed in tumors (Tubbs & Nussenzweig, 2017). We provide experimental evidence that Cyclin E upregulation is essential for tumor growth in this setting, identifying it as a central oncogenic driver in *Yki + dPGC1* knockdown tumors.

In sum, our findings demonstrate that *dPGC1* deregulation in cells with Yki activation leads to aberrant mitochondrial dynamics, increased Cyclin E levels, and DNA damage, factors that converge on key hallmarks of cancer, including altered cell proliferation, metabolism, and genomic instability. These results underscore the complexity of mitochondrial regulation in tumor biology and highlight the need to dissect the specific contributions of individual regulators of mitochondrial function. Targeting components of the mitochondrial fusion machinery may offer novel therapeutic strategies to limit Yki/YAP-driven tumor growth.

## Materials and Methods

### Drosophila strains

The following stocks used in this paper were obtained from the Bloomington *Drosophila* Stock Center: *dPGC1^1^* (synonym of *srl^KG08646^*) (14965), *hsFLP, UAS-GFP.U;; tub-Gal4 FRT82B, tub-Gal80* (86311), *FRT82B* (2051), *FRT82B Wts^X1^* (44251), *Gug-Gal4* (6773), *act-Gal4* (4414), *Mito-GFP* (8443), *yw* (1495), *UAS-LacZ* (3956), *UAS-dPGC1^RNAi^* (33914), *EP-dPGC1* (20009), *UAS-dMfn* (67157), *UAS-dMfn^RNAi^* (55189), *EP-Opa1* (20054), *UAS-Opa1^RNAi^* (32358), *UAS-miro* (51646), and *UAS-dap^RNAi^* (64026). The *UAS-dPGC1^shRNA^* (v330271) and *miro^RNAi^* (v106683) stocks were obtained from the Vienna *Drosophila* Stock Center. The *CycE^05306^* stock was a kind gift from Helena Richardson. The following stocks used in this study are described in the following references: *MS1096-Gal4* (Capdevila & Guerrero, 1994), *ap-Gal4 (Calleja et al, 1996)*, *UAS-Yki (Huang et al., 2005)*, *UAS-InR29.4* (*UAS-InR*) (Mirth *et al*, 2005), and *UAS-EGFR* (Buff *et al*, 1998).

*Drosophila* strains were raised on food that was made in-house containing agar (8g/L), brewer’s yeast (23.6g/L), dextrose (50.8g/L) and corn meal (58g/L). Depending on the experiment, flies were housed in humidity-controlled incubators set at either 18°C or 25°C with a 12/12-hour light dark cycle.

### List of Genotypes

Figure 1

(A, B) *yw* corresponds to *yw/yw*.

*dPGC1^1^ corresponds to dPGC1^1^/dPGC1^1^*.

(C-E) Control corresponds to *MS1096-Gal4, UAS-RFP/+; +/+; UAS-LacZ/+. MS1096>dPGC1^RNAi^* corresponds to *MS1096-Gal4, UAS-RFP/+; +/+; UAS-dPGC1^RNAi^/+*.

*MS1096>dPGC1* corresponds to *MS1096-Gal4, UAS-RFP/+; +/+; EP-dPGC1/+*.

Figure 2

(A-E; J-Q) Control corresponds to *ap-Gal4, UAS-CD8-GFP/+; tub-Gal80ts/UAS-LacZ. ap>dPGC1^RNAi^* corresponds to *ap-Gal4, UAS-CD8-GFP/+; tub-Gal80ts/UAS-dPGC1^RNAi^. ap>Yki* corresponds to *ap-Gal4, UAS-CD8-GFP/+; UAS-Yki, tub-Gal80ts/UAS-LacZ. ap>Yki+dPGC1^RNAi^* corresponds to *ap-Gal4, UAS-CD8-GFP/+; UAS-Yki, tub-Gal80ts/UAS-dPGC1^RNAi^*.

(F-I) Control corresponds to *hsFLP, UAS-GFP.U/+; +/+; tub-GAL4 FRT82B tub-GAL80/FRT82B Sb*.

*Wts^X1^* corresponds to *hsFLP, UAS-GFP.U/+; +/+; tub-Gal4 FRT82B tub-Gal80/FRT82B Wts^X1^*. *Wts^X1^+dPGC1^shRNA^* corresponds to *hsFLP, UAS-GFP.U/+; UAS-dPGC1^shRNA^/+; tub-Gal4 FRT82B tub-Gal80/ FRT82B Wts^X1^*.

Figure 3

A. The experimental condition corresponds to *ap-Gal4, UAS-CD8-GFP/+; UAS-Yki, tub-Gal80ts/UAS-dPGC1^RNAi^* and the control used for normalization corresponds to *ap-Gal4, UAS-CD8-GFP/+; UAS-Yki, tub-Gal80ts/UAS-LacZ*.
B. *ap-Gal4/+; UAS-Mito-GFP/+*.
C. *Gug-Gal4, UAS-Mito-GFP/+*.
D. *Gug-Gal4, UAS-Mito-GFP/UAS-dMfn*.
E. *UAS-dMfn^RNAi^/+; Gug-Gal4, UAS-Mito-GFP/+*.
F. *EP-Opa1/+; Gug-Gal4, UAS-Mito-GFP/+*.
G. *Gug-Gal4, UAS-Mito-GFP/UAS-Opa1^RNAi^*.
H. *UAS-miro/+; Gug-Gal4, UAS-Mito-GFP/+*.
I. *UAS-miro^RNAi^/+; Gug-Gal4, UAS-Mito-GFP/+*.

Figure 4

*ap>Yki* corresponds to *ap-Gal4, UAS-CD8-GFP/+; UAS-Yki, tub-Gal80ts/UAS-LacZ. ap>Yki+dPGC1^RNAi^* corresponds to *ap-Gal4, UAS-CD8-GFP/+; UAS-Yki, tub-Gal80ts/UAS-dPGC1^RNAi^*.

*ap>Yki+dPGC1^RNAi^+dMfn^RNAi^*corresponds to *ap-Gal4, UAS-CD8-GFP/UAS-dMfn^RNAi^; UAS-Yki, tub-Gal80ts/UAS-dPGC1^RNAi^*.

Figure 5

Control corresponds to *ap-Gal4, UAS-CD8-GFP/+; tub-Gal80ts/UAS-LacZ*.

*ap>Yki* corresponds to *ap-Gal4, UAS-CD8-GFP/+; UAS-Yki, tub-Gal80ts/UAS-LacZ. ap>Yki+dPGC1^RNAi^* corresponds to *ap-Gal4, UAS-CD8-GFP/+; UAS-Yki, tub-Gal80ts/UAS-dPGC1^RNAi^*.

*ap>Yki+dPGC1^RNAi^+dMfn^RNAi^*corresponds to *ap-Gal4, UAS-CD8-GFP/UAS-dMfn^RNAi^; UAS-Yki, tub-Gal80ts/UAS-dPGC1^RNAi^*.

*ap>Yki+dPGC1^shRNA^*corresponds to *ap-Gal4, UAS-CD8-GFP/UAS-dPGC1^shRNA^; UAS-Yki, tub-Gal80ts/+*.

*ap>Yki+dPGC1^shRNA^+Opa1^RNAi^*corresponds to *ap-Gal4, UAS-CD8-GFP/UAS-dPGC1^shRNA^; UAS-Yki, tub-Gal80ts/UAS-Opa1^RNAi^*.

*ap>Yki+dMfn* corresponds to *ap-Gal4, UAS-CD8-GFP/+; UAS-Yki, tub-Gal80ts/UAS-dMfn. ap>Yki+Opa1* corresponds to *ap-Gal4, UAS-CD8-GFP/EP-Opa1; UAS-Yki, tub-Gal80ts/+*.

Figure 6

Control corresponds to *ap-Gal4, UAS-CD8-GFP/+; tub-Gal80ts/UAS-LacZ*.

*ap>; CycE^05306^* corresponds to *ap-Gal4, UAS-CD8-GFP/CycE^05306^; tub-Gal80ts/+. ap>Yki* corresponds to *ap-Gal4, UAS-CD8-GFP/+; UAS-Yki, tub-Gal80ts/UAS-LacZ*.

*ap>Yki+dap^RNAi^* corresponds to *ap-Gal4, UAS-CD8-GFP/UAS-dap^RNAi^; UAS-Yki, tub-Gal80ts/+. ap>Yki+dPGC1^RNAi^* corresponds to *ap-Gal4, UAS-CD8-GFP/+; UAS-Yki, tub-Gal80ts/UAS-dPGC1^RNAi^*.

*ap>Yki* corresponds to *ap-Gal4, UAS-CD8-GFP/+; UAS-Yki, tub-Gal80ts/UAS-LacZ. ap>Yki+dap^RNAi^* corresponds to *ap-Gal4, UAS-CD8-GFP/UAS-dap^RNAi^; UAS-Yki, tub-Gal80ts/+*.

*ap>Yki+dPGC1^RNAi^; CycE^05306^* corresponds to *ap-Gal4, UAS-CD8-GFP/CycE^05306^; UAS-Yki, tub-Gal80ts/UAS-dPGC1^RNAi^*.

Figure S1

*yw* corresponds to *yw/yw*.

*dPGC1^1^ corresponds to dPGC1^1^/dPGC1^1^*.

Figure S2

*act>* corresponds to *act-Gal4/+*.

*act>dPGC1^RNAi^* corresponds to *act-Gal4/UAS-dPGC1^RNAi^. act>dPGC1^shRNA^* corresponds to *dPGC1^shRNA^/+; act-Gal4/+. act>dMfn* corresponds to *act-Gal4/UAS-dMfn. act>dMfn^RNAi^*corresponds to *UAS-dMfn^RNAi^/+; act-Gal4/+. act>Opa1* corresponds to *EP-Opa1/+; act-Gal4/+. act>Opa1^RNAi^* corresponds to *act-Gal4/UAS-Opa1^RNAi^. act>miro* corresponds to *UAS-miro/+; act-Gal4/+. act>miro^RNAi^*corresponds to *UAS-miro^RNAi^/+; act-Gal4/+*.

Figure S3

Control corresponds to *ap-Gal4, UAS-CD8-GFP/+; tub-Gal80ts/UAS-LacZ. ap>dPGC1^RNAi^* corresponds to *ap-Gal4, UAS-CD8-GFP/+; tub-Gal80ts/UAS-dPGC1^RNAi^*.

Figure S4

(A-C) *ap>EGFR* corresponds to *ap-Gal4, UAS-CD8-GFP/+; UAS-EGFR, tub-Gal80ts/+. ap>EGFR+dPGC1^RNAi^*corresponds to *ap-Gal4, UAS-CD8-GFP/+; UAS-EGFR, tub-Gal80ts/UAS-dPGC1^RNAi^*.

(D-F) *ap>InR* corresponds to *ap-Gal4, tub-Gal80ts/+; UAS-InR/+*.

*ap>InR+dPGC1^RNAi^*corresponds to *ap-Gal4, tub-Gal80ts/+; UAS-InR/UAS-dPGC1^RNAi^*.

Figure S5

Control corresponds to *ap-Gal4, UAS-CD8-GFP/+; tub-Gal80ts/UAS-LacZ*.

*ap>dPGC1^shRNA^* corresponds to *ap-Gal4, UAS-CD8-GFP/ UAS-dPGC1^shRNA^; tub-Gal80ts/+. ap>Yki* corresponds to *ap-Gal4, UAS-CD8-GFP/+; UAS-Yki, tub-Gal80ts/UAS-LacZ. ap>Yki+dPGC1^shRNA^* corresponds to *ap-Gal4, UAS-CD8-GFP/UAS-dPGC1^shRNA^; UAS-Yki, tub-Gal80ts/+*.

Fig S6

*ap>Yki* corresponds to *ap-Gal4, UAS-CD8-GFP/+; UAS-Yki, tub-Gal80ts/UAS-LacZ. ap>Yki+dPGC1^RNAi^* corresponds to *ap-Gal4, UAS-CD8-GFP/+; UAS-Yki, tub-Gal80ts/UAS-dPGC1^RNAi^*.

Fig S7

The experimental condition corresponds to *ap-Gal4, UAS-CD8-GFP/+; tub-Gal80ts/UAS-dPGC1^RNAi^*and the control used for normalization corresponds to *ap-Gal4, UAS-CD8-GFP/+; tub-Gal80ts/UAS-LacZ*.

Fig S8

*ap>Yki* corresponds to *ap-Gal4, UAS-CD8-GFP/+; UAS-Yki, tub-Gal80ts/UAS-LacZ. ap>Yki+dPGC1* corresponds to *ap-Gal4, UAS-CD8-GFP/+; UAS-Yki, tub-Gal80ts/EP-dPGC1*.

Figure S9

Control corresponds to *Gug-Gal4, UAS-Mito-GFP/+*.

*Gug>dMfn* corresponds to *Gug-Gal4, UAS-Mito-GFP/UAS-dMfn. Gug>Mfn^RNAi^* corresponds to *UAS-dMfn^RNAi^/+; Gug-Gal4, UAS-Mito-GFP/+. Gug>Opa1* corresponds to *EP-Opa1/+; Gug-Gal4, UAS-Mito-GFP/+*.

*Gug>Opa1^RNAi^*corresponds to *Gug-Gal4, UAS-Mito-GFP/UAS-Opa1^RNAi^*.

*Gug>miro* corresponds to *UAS-miro/+; Gug-Gal4, UAS-Mito-GFP/+. Gug>miro^RNAi^* corresponds to *UAS-miro^RNAi^/+; Gug-Gal4, UAS-Mito-GFP/+*.

Figure S10

Control corresponds to *ap-Gal4, UAS-CD8-GFP/+; tub-Gal80ts/UAS-LacZ. ap>Mfn* corresponds to *ap-Gal4, UAS-CD8-GFP/+; tub-Gal80ts/ UAS-dMfn. ap>Mfn^RNAi^* corresponds to *ap-Gal4, UAS-CD8-GFP/ UAS-dMfn^RNAi^; tub-Gal80ts/+. ap>Opa1* corresponds to *ap-Gal4, UAS-CD8-GFP/EP-Opa1; tub-Gal80ts/+*.

*ap>Opa1^RNAi^*corresponds to *ap-Gal4, UAS-CD8-GFP/+; tub-Gal80ts/ UAS-Opa1^RNAi^*.

Figure S11

(A-C) *ap>Yki+dP*GC1*^RNAi^*corresponds to *ap-Gal4, UAS-CD8-GFP/+; UAS-Yki, tub-Gal80ts/UAS-dP*GC1*^RNAi^*.

*ap>Yki+dP*GC1*^RNAi^+miro^RNAi^*corresponds to *ap-Gal4, UAS-CD8-GFP/UAS-miro^RNAi^; UAS-Yki, tub-Gal80ts/UAS-dP*GC1*^RNAi^*.

(D-F) *ap>Yki* corresponds to *ap-Gal4, UAS-CD8-GFP/+; UAS-Yki, tub-Gal80ts/UAS-LacZ. ap>Yki+miro* corresponds to *ap-Gal4, UAS-CD8-GFP/UAS-miro; UAS-Yki, tub-Gal80ts/+*.

Figure S12

*ap>Yki* corresponds to *ap-Gal4, UAS-CD8-GFP/+; UAS-Yki, tub-Gal80ts/UAS-LacZ. ap>Yki+dPGC1^RNAi^* corresponds to *ap-Gal4, UAS-CD8-GFP/+; UAS-Yki, tub-Gal80ts/UAS-dPGC1^RNAi^*.

Figure S13

Figure S14

*yw* corresponds to *yw/yw*.

*dPGC1^1^ corresponds to dPGC1^1^/dPGC1^1^*.

### Temporal Control of Transgene Expression by Gal80^ts^

For experiments using ap-Gal4 or act-Gal4, the Gal4/Gal80^ts^ system was used to prevent lethality during early development. This system allowed UAS-driven transgenes to be controlled in a temperature-dependent manner. *Drosophila* crosses were kept for 2 days at 18°C to lay eggs. After 2 days, crosses were flipped into new vials and the old vials containing eggs were kept for 2 additional days at 18°C before being transferred to a 29°C incubator for transgene expression to be induced. To ensure that wing discs were analyzed at the same developmental stage, larvae with hyperplastic tumors were dissected after 5 days at 29°C, whereas giant larvae containing neoplastic tumors were dissected after 7 days at 29°C.

### Induction of Mitotic Clones

Mitotic recombination clones were generated using the FLP/FRT system, employing the MARCM (Mosaic Analysis with a Repressible Cell Marker) technique. Larvae carrying the necessary transgenes (see the list of genotypes for details) were maintained at 25°C for 48–96 hours after egg laying. To induce FLP recombinase expression, vials were heat-shocked by immersion in a circulating water bath at 37°C for 60 minutes. Following heat shock, larvae were returned to 25°C and allowed to develop for an additional 96 hours to permit clone growth. Larvae were dissected at the third instar wandering stage, corresponding to 96 hours after heat shock, for analysis of imaginal discs.

### Immunohistochemistry

Larvae were dissected in PBS and fixed in 3.7% formaldehyde / PBS for 20 minutes at room temperature. Samples were then washed in 0.2% Triton / PBS (PBT) and blocked for an hour at room temperature in blocking buffer (BBT) made of 0.2% Triton, 5mM NaCl, and 3% BSA diluted in PBS. Primary antibodies were diluted in BBT, and the samples were left in primary antibody solution overnight at room temperature. Longer primary antibody incubations were done at 4°C. After removing the primary antibody and performing washes with BBT, samples were incubated for 2 hours at room temperature with the relevant fluorescent secondary antibodies and DAPI diluted in BBT. After removing the secondary antibody and performing washes with PBT, samples were mounted in 90% glycerol / PBS containing 0.5% N-propyl gallate. Samples were imaged with the Leica SP8 confocal microscope. For stainings intended for mitochondrial analysis, BBT and PBT contained 0.2% Tween instead of 0.2% Triton.

The following primary antibodies were used: mouse anti-MMP1 (Developmental Studies Hybridoma Bank, 3A6B4/5H7B11/3B8D12 mixed in equal volumes), rabbit anti-pH2Av (Rockland, 600-401-914, dilution 1:1000), rabbit anti-PH3 (Cell Signal Technology, 9701, dilution 1:100), and rabbit anti-Cyclin E (Santa Cruz, 33748, dilution 1:100). The following secondary antibodies were used: Alexa Fluor 635-phalloidin to stain F-actin (Invitrogen, A34054, dilution 1:200), anti-mouse Alexa Fluor 555 (Invitrogen, A21425, dilution 1:200), and anti-rabbit Alexa Fluor 555 (Invitrogen, A21430, dilution 1:200). DAPI (Invitrogen, D1306) was used at a 600 nM concentration to stain the nuclei.

### Analysis of Mitochondrial Membrane Potential

Wing discs from *Drosophila* larvae were dissected in Schneider’s medium (Sigma, Ref: S9895). After dissection, wing discs were incubated in 100 nM TMRE (Sigma, Ref: 87917, dissolved in Schneider’s medium) for 20 minutes with agitation. Then, wing discs were washed once with Schneider’s medium for 5 minutes, rinsed with PBS, and mounted in Schneider’s medium. Samples were immediately imaged after mounting with the Leica SP8 confocal microscope. All steps were performed at room temperature.

### Electron Microscopy

Larval wing discs were dissected in PBS and fixed in 2% glutaraldehyde / 0.1M sodium cacodylate buffer at room temperature. Sample preparation was performed by the Core Facility for Integrative Microscopy at the University of Copenhagen, where tissues were fixed, embedded, stained, dehydrated, infiltrated, embedded, and cut into thin sections for imaging. All specimens were imaged at the Core Facility for Integrative Microscopy with a Transmission Electron Microscope (Philips CM100, FEL). Digital images were recorded with an OSIS Veleta digital slow-scan 2k x 2k CCD camera and the ITEM software package. Mitochondrial morphologies reflected in these 2D micrographs were analyzed with Fiji software. Mitochondria were manually outlined and quantified for area, perimeter, and aspect ratio (calculated as MaxFeret/MinFeret to assess elongation). At least 164 mitochondria were quantified per genotype.

### RNA Extraction, cDNA Synthesis, and qPCR

For experiments performed with ap-Gal4, RNA was extracted from wing discs dissected from third instar wandering larvae with the TRIzol reagent (Life Technologies, Ref: 15596026). For experiments performed with act-Gal4, RNA was extracted from whole third instar wandering larvae with the RNeasy Mini Kit (Qiagen, Ref: 74106). Total RNA was treated with RQ1 DNase I (Promega, Ref: M6101) and converted to cDNA using the SuperScript III Reverse Transcriptase kit (Life technologies, Ref: 18080-044). qPCR was performed using 5x HOT FIREPol EvaGreen qPCR Mix Plus (Solis Biodyne, Ref: 08-24-00001) on the QuantStudio 6 Flex Real-Time PCR machine (Applied Biosystems). The primers used are listed below.

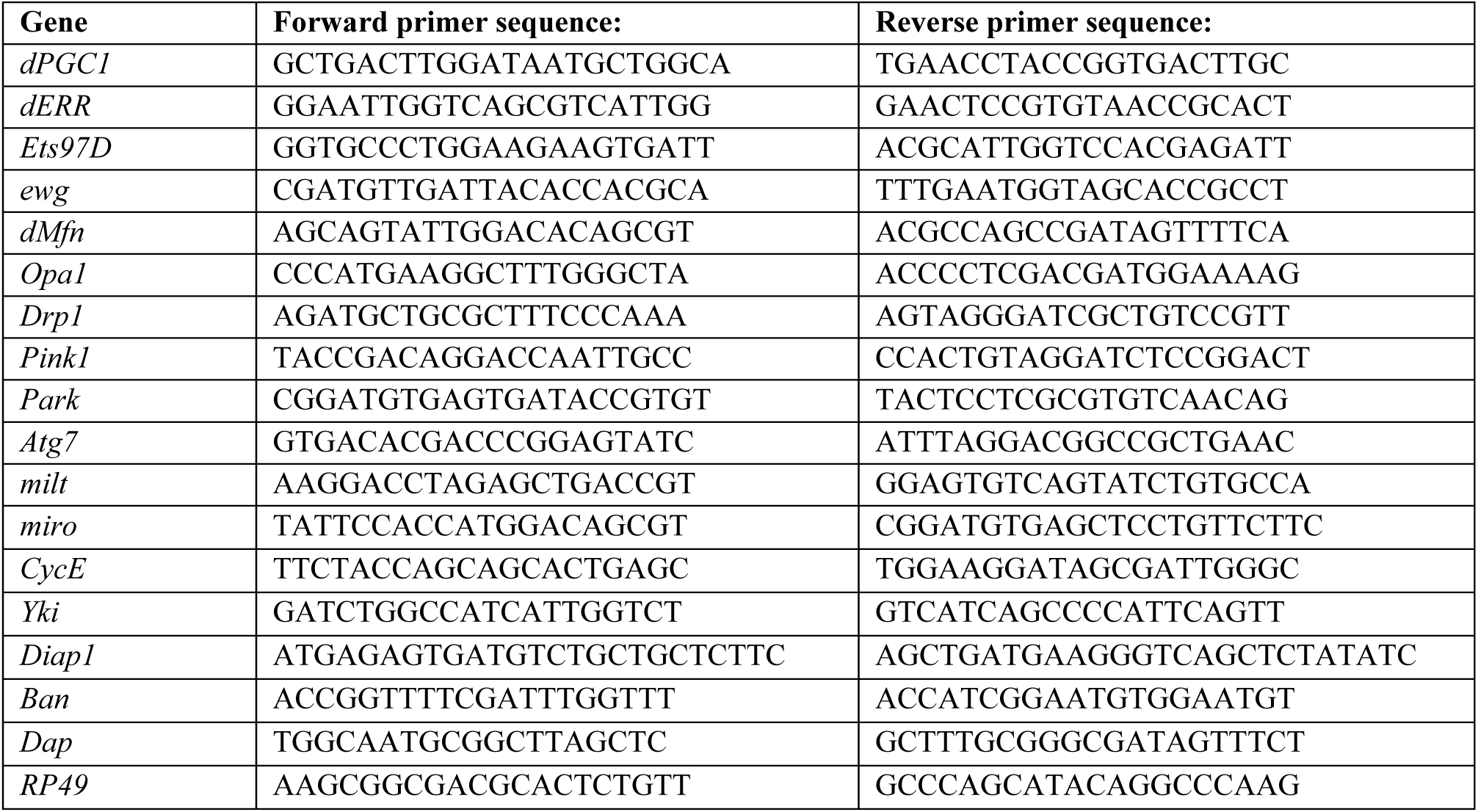

### Western Blot

Wing discs were dissected from *Drosophila* larvae and transferred into RIPA protein extraction and immunoprecipitation buffer (Sigma-Aldrich, Ref: R0278) containing cOmplete Protease Inhibitor Cocktail (Merck, Ref: 04693124001), PhosSTOP (Merck, Ref: 04906837001), and 100mM PMSF (Sigma-Aldrich, Ref: 93482). Wing discs were homogenized by pipetting up and down and then leaving samples on ice for 30 minutes. Afterwards, samples were sonicated three times for three cycles with the following settings: 100% amplitude, 1 cycle. Samples were then centrifuged for 20 minutes at 15000g and 4°C to obtain the supernatant that was used for western blots. Protein concentration was quantified using the Pierce BCA protein assay kit (Thermo Fisher Scientific, Ref: 23227). Western blots were performed using 15 μg of protein per sample. Western blot membranes were blocked for an hour at room temperature with 5% skimmed milk (Fluka, Ref: 70166-500G) dissolved in TBST. After washing with TBST, the membranes were incubated overnight at 4°C with primary antibodies diluted in TBST. After removing the primary antibody and performing washes with TBST, samples were incubated for 1 hour at room temperature with the relevant HRP-conjugated secondary antibodies. After removing the secondary antibodies and washing with TBST, the Pierce ECL Western Blotting Substrate kit (Thermo Fisher Scientific, Ref: 32106) was used to visualize protein levels on the Amersham Imager 600 (GE Healthcare).

The following primary antibodies were used: rabbit anti-Cyclin E (Santa Cruz, 33748, dilution 1:1000) and mouse anti-α-Tubulin (Developmental Studies Hybridoma Bank, 12G10, dilution 1:1000). The following HRP-conjugated secondary antibodies were used: polyclonal goat anti-rabbit-HRP (Invitrogen, 31460, dilution 1:10000) and polyclonal goat anti-mouse-HRP (Dako, P0447, dilution 1:5000). Note that since Cyclin E and α-Tubulin have similar molecular weights, membranes had to be stripped to detect both proteins without any crossover.

### Image Analysis and Processing

Image analyses were performed using Fiji (ImageJ) software. Specific quantification procedures are described in the sections below. Final figures were assembled using Adobe Photoshop.

### Quantification of *Drosophila* Wing Imaginal Disc and Clone Size

To quantify wing imaginal disc size, the “threshold” function was first used to select either the GFP-positive area or the DAPI-positive area, depending on the experiment. After setting a threshold, areas were determined using the “analyze particles” option. A minimum of 10 discs were quantified per genotype. To quantify clone size, individual clones in each wing disc were selected with the “polygon” tool, and their area was directly measured using the “measure” option. For each genotype, at least 65 clones were quantified from a total of a minimum of 10 discs. For both wing disc and clone size quantifications, results were expressed as the fold change of GFP or DAPI area relative to the corresponding control.

### Quantification of *Drosophila* Adult Wing Size

Crosses were allowed to lay eggs on apple agar plates with yeast paste at 25°C for 10 to 14 hours. Once eggs hatched and L1 larvae emerged, 20 larvae were transferred to vials containing standard *Drosophila* food, and these vials were kept at 25°C until adult flies eclosed. Wings were fixed by introducing adult flies into a 25% glycerol / 75% ethanol mixture for at least 24 hours. After that, wings were plucked and mounted in glycerol for imaging. Images of adult *Drosophila* wings were taken with a Leica M165 stereomicroscope.

To measure adult wing area, the “polygon” selection tool in Fiji was used to draw a border around the wing edges. Next, the “measure” option was used to quantify the wing area. Results were expressed as the fold change in wing area relative to the control. A total of 20 wings were quantified per genotype.

### Quantification of Mitochondrial Shape in the Peripodial Membrane

The Gug-Gal4 driver was used to drive the expression of the *Mito-GFP* mitochondria marker in the peripodial membrane. After obtaining images of peripodial membrane cells, two filters were applied to the GFP channel, unsharp mask (radius = 10.0 pixels, mask strength = 0.9) and median filtering (radius = 3), to enhance and smooth the Mito-GFP signal, respectively (Terriente-Felix *et al*, 2020). Next, the “threshold” function was used to create a binary mask, and the “Mitochondrial Analyzer 2D Analysis” plugin from ImageJ (Chaudhry *et al*, 2020) was used to calculate the mean form factor. Mitochondria from a minimum of 15 images were quantified per genotype.

### Quantification of TMRE Intensity

For each image, the area of the wing disc containing mitochondria stained with TMRE was selected using the “threshold” function on the TMRE channel. The mean gray value of only the thresholded selection was obtained with the “measure” function. Results were expressed as the fold change of TMRE intensity relative to the control. Importantly, comparisons were only made between wing discs that had been stained and imaged together, meaning that fold changes were always calculated considering the mean of the control for each individual experiment and then all fold changes were pooled for the statistical analysis. For each genotype, a minimum of 23 wing discs from a total of 5 experiments were quantified.

### Quantification of Mmp1 Area

For each wing disc, the GFP-positive area was first measured as explained above and the GFP selection was added to the ROI manager. Afterwards, a “gaussian blur” filter of 1 was applied to the Mmp1 channel and the “threshold” function was used to select only the Mmp1-positive signal. The thresholded Mmp1 area within the previously selected GFP area was measured with the “measure” option within the ROI manager. Results were expressed as the Mmp1 area normalized to the GFP-positive area. A minimum of 11 wing discs were quantified per genotype.

### Quantification of F-actin and Cyclin E Intensity

Quantification of F-actin and Cyclin E intensities were done with the same procedure. The “polygon” tool was first used to select a DAPI-positive and GFP-positive (if applicable) area within the image. Then, the mean gray value of the corresponding channel (either F-actin or Cyclin E) within the selected area was obtained with the “measure” function. Results were expressed as the direct values of mean intensity (for F-actin) or as the fold change in mean intensity relative to the control (for Cyclin E). A minimum of 19 wing discs (for F-actin) or 13 wing discs (for Cyclin E) were quantified per genotype.

### Quantification of PH3-Positive Cells

First, the “polygon” tool was used to select a DAPI-positive and GFP-positive area, which was then measured with the “measure” function. Then, a “gaussian blur” filter of 3 was applied to the PH3 channel. Finally, the number of PH3-positive cells was measured with the “find maxima” tool. Results were expressed as the number of PH3-positive cells per GFP-positive area. A minimum of 19 wing discs were quantified per genotype.

### Quantification of pH2Av-Positive Foci

Maximum projections of Z-stack images of wing discs were used for this quantification. First, the GFP-positive area was measured as previously described. Then, the number of pH2Av-positive foci was measured using the “3D Objects Counter”. Results were expressed as the fold change in number of pH2Av-positive foci per area relative to the control. A minimum of 18 wing discs were quantified per genotype.

### Quantification of Protein Levels in Western Blot Membranes

Western blot membranes were loaded to Fiji, and the “rectangle” tool was used to select the corresponding bands, fitting all genotypes in the same rectangle. Within the “gels” function, “select first lane” was used to set this rectangle as a lane, and then “plot lanes” was used to generate a lane profile plot showcasing intensity peaks corresponding to each band within the rectangle. The “line” tool was used to draw base lines for each peak, generating closed areas that did not include the background signal, and these peak areas were then measured. For each genotype, areas obtained for Cyclin E bands were normalized to the corresponding α-Tubulin bands. Results were expressed as the fold change in Cyclin E/α-Tubulin peak areas for each membrane, and then all fold changes were pooled for the statistical analysis. A total of 7 western blot membranes were quantified.

### Statistics

Graphs and statistical analyses were done using GraphPad Prism 10. Data is always presented as full-range box plots. The number of biological replicates quantified per sample and the statistical tests applied to determine statistical significance are indicated in the corresponding figure legends.

## Acknowledgements

We thank Helena Richardson, Kim F. Rewitz, and Takashi Koyama for reagents. We thank Bloomington *Drosophila* Stock Center and Vienna *Drosophila* Resource Center for fly stocks; Developmental Studies Hybridoma Bank for antibodies; and the fly community for sharing reagents. We also thank the Center for Integrated Microscopy (CFIM) at the University of Copenhagen for electron microscopy support.

## Conflict of interest

The authors declare that the research was conducted in the absence of any commercial or financial relationships that could be construed as a potential conflict of interest.

## Funding

This work was supported by the Independent Research Fund Denmark Grant 0134-00045B, Novo Nordisk Foundation Grant NNF18OC0052223, a grant from the Neye Foundation, and a grant from Grosserer Alfred Nielsen og Hustru’s fond.

## Notes

### Competing Interest Statement

The authors have declared no competing interest.

### Summary of Updates

We have undertaken a substantial revision that includes new experimental data, refined analyses, and clearer presentation of our findings. Specifically, we have addressed concerns about RNAi efficiency and protein-level validation, expanded our genetic models to include loss-of-function contexts, and clarified the interpretation of mitochondrial morphology using both confocal and electron microscopy. We also incorporated new data on Cyclin E regulation and mitochondrial membrane potential to strengthen the mechanistic link between dPGC1 depletion and Yki-driven tumorigenesis. These revisions enhance the coherence and impact of the study. We are confident that the revised manuscript presents a more robust and compelling case for the role of dPGC1 as a context-dependent tumor suppressor and that it will be of broad interest to the fields of developmental biology, cancer metabolism, and mitochondrial dynamics.

