## Supplementary Figures for "Control of mitochondrial dynamics by dPGC1 limits Yorkie-induced oncogenic growth in *Drosophila*"

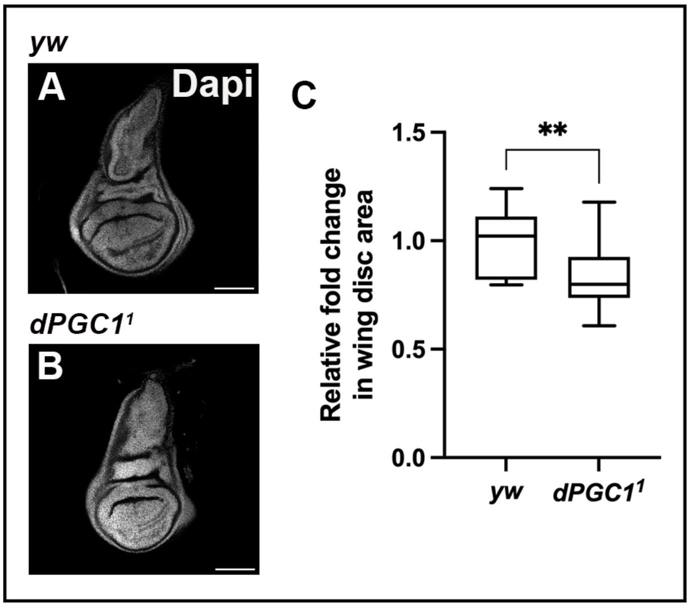


**Figure S1. *dPGC1* Mutant Wing Discs**

**(A, B)** Confocal micrographs of *Drosophila* wing imaginal discs from the following genotypes: *yw/yw* (control, A) and *dPGC1$^1$/dPGC1$^1$* (mutant, B). DAPI labels the DNA and is shown in grayscale. Scale bars, 100 µm.

**(C)** Quantification of wing disc area of the genotypes in A and B. Wing disc size was normalized to the mean area of the control (*yw/yw*). Statistical significance was determined using an unpaired t-test (n = 16 [*yw*], n = 13 [*dPGC1$^1$*]). $^{**}p < 0.01$.

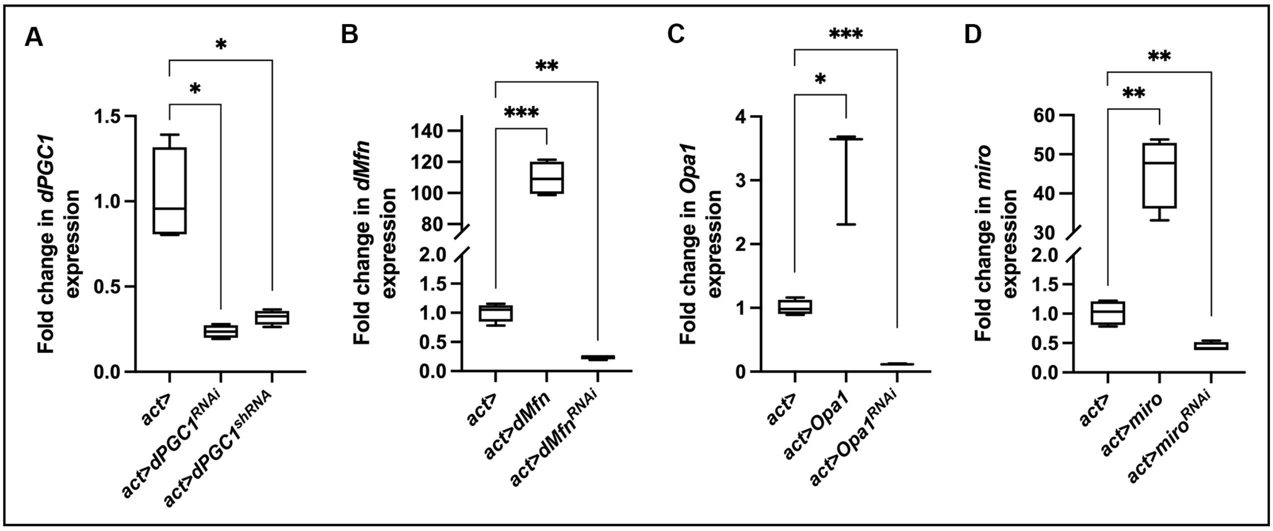


**Figure S2. UAS-Driven Transgene Efficiency**

mRNA quantification by qPCR of the transgenes used to modulate the genes *dPGC1* (A), *dMfn* (B), *Opa1* (C), and *miro* (D). The transgenes were expressed under the control of the act-Gal4 driver and whole larvae were used for the analysis. The control genotype is *act-Gal4/+*. Each genotype was run in three to five biological replicates. RP49 was used as the housekeeping gene. Statistical significance was determined using unpaired t-tests with or without Welch's correction depending on whether variances were significantly different or not, respectively. *p < 0.05, **p < 0.01, ***p < 0.001.

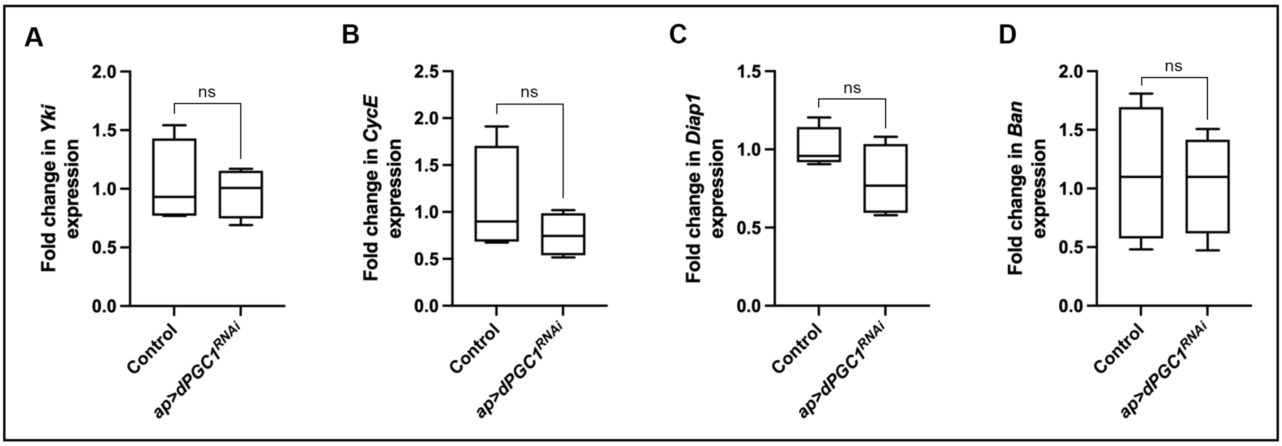


**Figure S3. *Yki* Expression and Activity in Discs with Reduced dPGC1**

mRNA quantification by qPCR of *Yki* (A) and the Yki target genes *CycE* (B), *Diap1* (C), and *Ban* (D) in *ap-Gal4, UAS-GFP, UAS-dPGC1-RNAi* wing imaginal discs. The control genotype is *ap-Gal4, UAS-GFP, UAS-LacZ*. Each genotype was run in four biological replicates. RP49 was used as the housekeeping gene. Statistical significance was determined using unpaired t-tests. ns, non-significant.

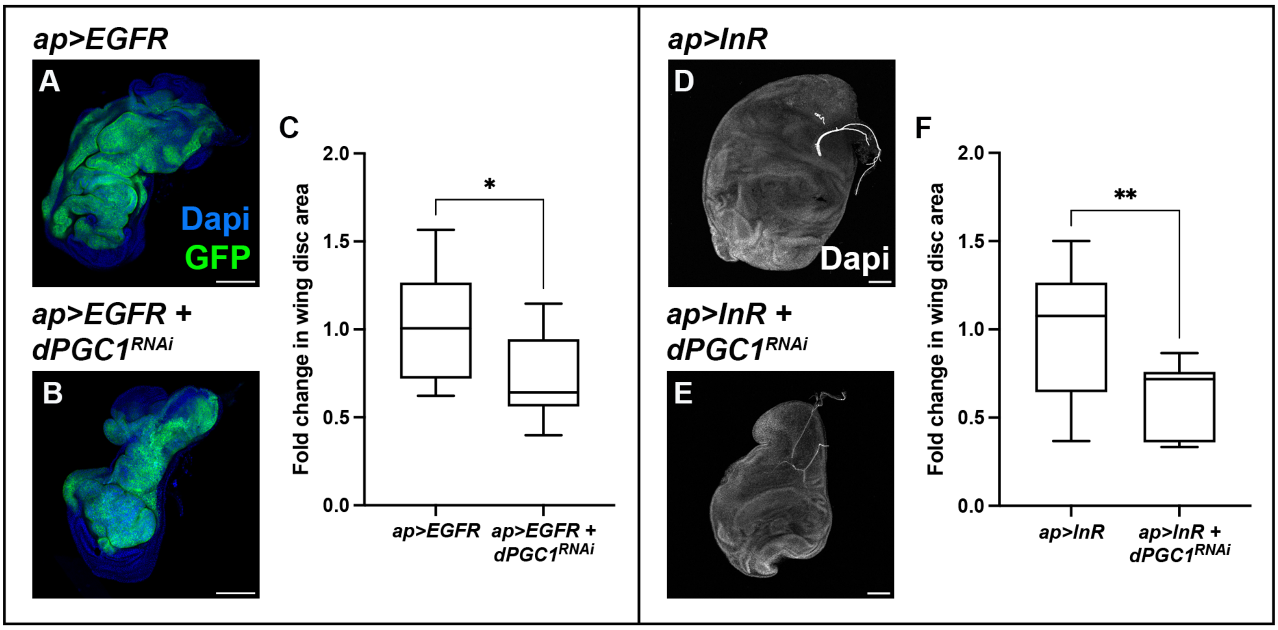


**Figure S4. *dPGC1* Depletion Does Not Cooperate with the oncogenes *EGFR* and *InR***

**(A, B)** Confocal micrographs of wing imaginal discs of the following genotypes: *ap-Gal4, UAS-EGFR, UAS-GFP, UAS-LacZ* (A) and *ap-Gal4, UAS-EGFR, UAS-GFP, UAS-dPGC1-RNAi* (B). GFP is shown in green. DAPI labels the DNA and is shown in blue. Scale bars, 100 µm.

**(C)** Quantification of wing disc area (GFP-positive area) of the genotypes in A and B. GFP-positive area was normalized to the mean of the control (*ap>EGFR*). Statistical significance was determined using an unpaired t-test (n = 11 [*ap>EGFR*], n = 11 [*ap>EGFR*, *dPGC1-RNAi*]). *$p < 0.05$.

**(D, E)** Confocal micrographs of wing imaginal discs of the following genotypes: *ap-Gal4, UAS-InR, UAS-GFP* (GFP not shown) (D) and *ap-Gal4, UAS-InR, UAS-dPGC1-RNAi* (E). DAPI labels the DNA and is shown in grayscale. Scale bars, 100 µm.

**(F)** Quantification of wing disc area (DAPI-positive area) of the genotypes in D and E. DAPI-positive area was normalized to the mean of the control (*ap>InR*). Statistical significance was determined using an unpaired t-test (n = 19 [*ap>InR*], n = 11 [*ap>InR, dPGC1-RNAi*]). **$p < 0.01$.

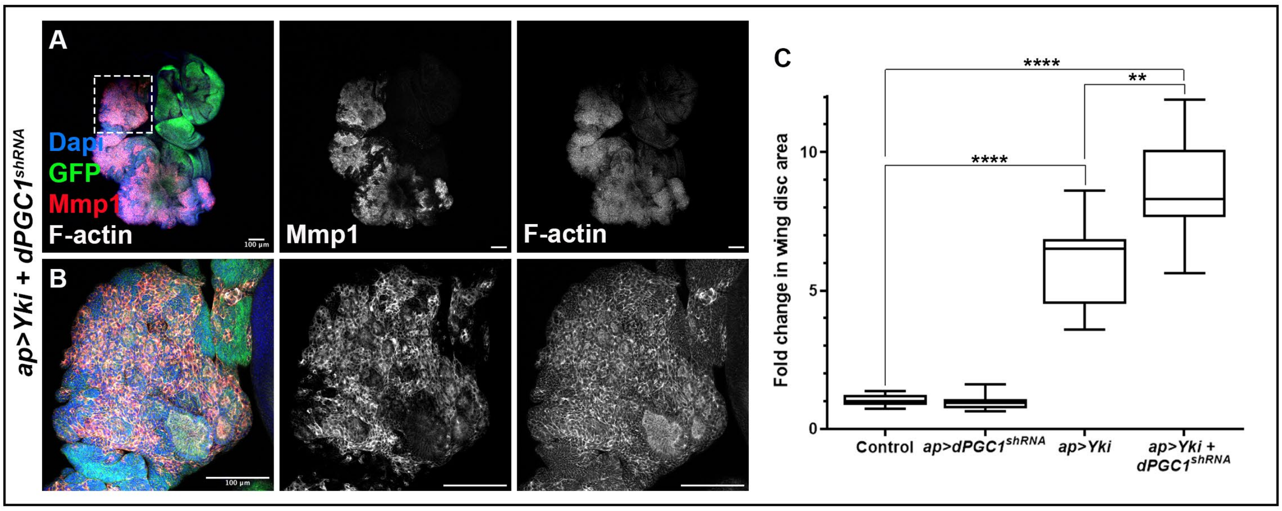


**Figure S5. Oncogenic Cooperation Between Yki and dPGC1**

**(A, B)** Confocal micrograph showing an *ap-Gal4, UAS-GFP, UAS-Yki, UAS-dPGC1-shRNA* third instar wing imaginal disc. F-actin labels cell polarity and is shown in grayscale. Mmp1 is shown in red. GFP is shown in green. DAPI labels the DNA and is shown in blue. The dashed white box in A indicates the region of the wing disc that is shown as a magnification in B. Scale bars, 100 µm.

**(C)** Quantification of GFP-positive area in third instar wing imaginal discs of the indicated genotypes. GFP-positive areas were normalized to the mean of the control (*ap-Gal4, UAS-GFP, UAS-LacZ*). Statistical significance was determined using unpaired t-test with Welch's correction (n = 10 [Control], n = 10 [*ap>dPGC1-shRNA*], n = 10 [*ap>Yki*], n = 10 [*ap>Yki, dPGC1-shRNA*]). **p < 0.01, ****p < 0.0001.

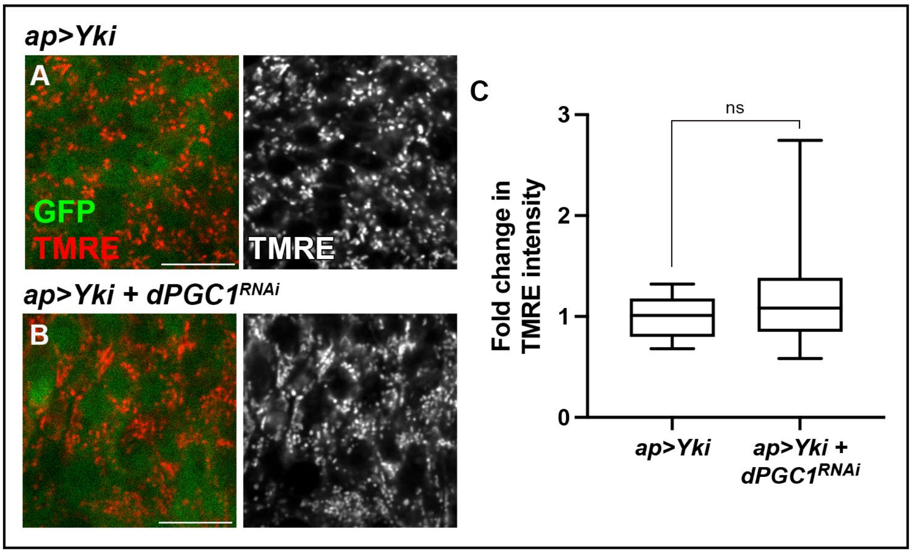


**Figure S6. Membrane Potential in *Yki+dPGC1-RNAi* tumors**

**(A, B)** Confocal micrographs showing magnifications from tumorous wing discs with the following genotypes: *ap-Gal4, UAS-Yki, UAS-GFP, UAS-LacZ* (A) and *ap-Gal4, UAS-Yki, UAS-GFP, UAS-dPGC1-RNAi* (B). GFP is shown in green. TMRE staining is shown in red. Scale bars, 10 µm.

**(C)** Quantification of TMRE intensity in the genotypes in A and B. TMRE intensity was normalized to the mean of the control (*ap>Yki*). Statistical significance was determined using a Mann-Whitney test for non-parametric data (n = 23 [*ap>Yki*], n = 56 [*ap>Yki, dPGC1-RNAi*). ns, non-significant.

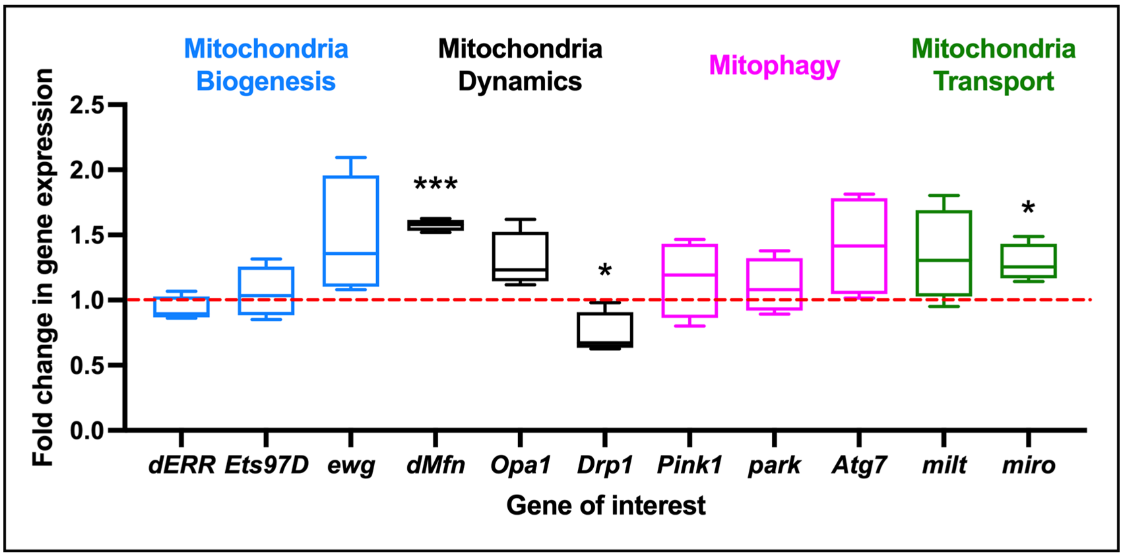


**Figure S7. Expression of Genes Controlling Mitochondrial Dynamics in Discs with Reduced dPGC1**

mRNA quantification by qPCR of the indicated genes in *ap-Gal4, UAS-GFP, UAS-dPGC1-RNAi* wing imaginal discs using *ap-Gal4, UAS-GFP, UAS-LacZ* as the control genotype. Genes are separated in different categories: mitochondria biogenesis (blue), mitochondria dynamics (black), mitophagy (pink), and mitochondria transport (green). Each genotype was run in four biological replicates. RP49 was used as the housekeeping gene. Statistical significance was determined using unpaired t-tests with Welch's correction for parametric data and Mann-Whitney tests for non-parametric data. *p < 0.05, ***p < 0.001.

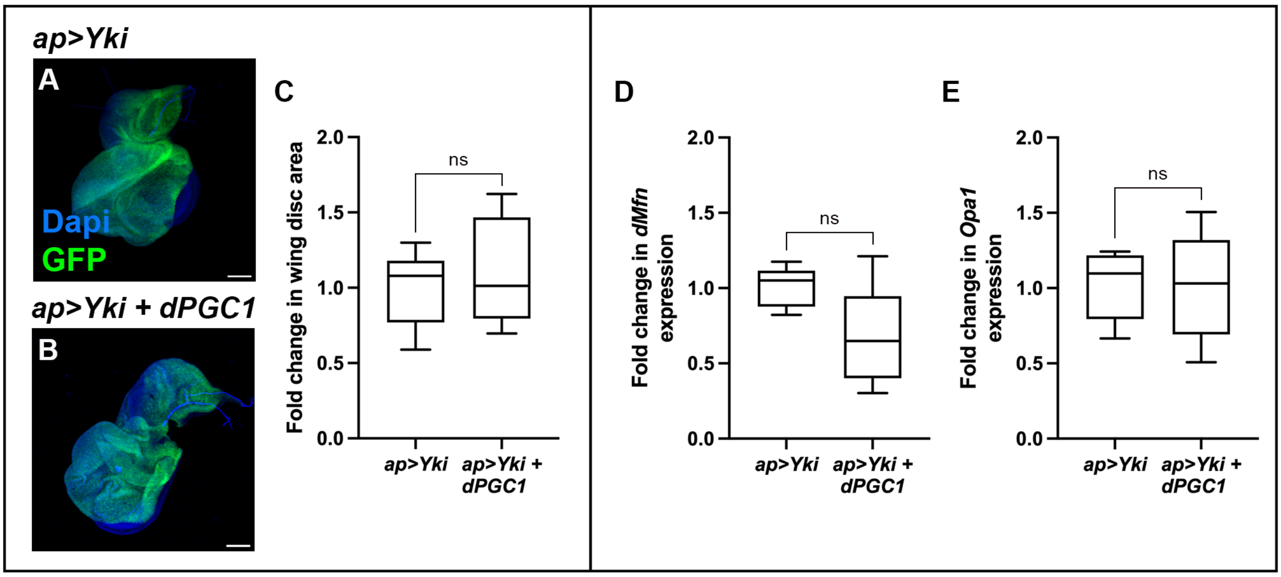


**Fig S8. *dPGC1* Overexpression in a Context of *Yki* Upregulation**

**(A, B)** Confocal micrographs of wing imaginal discs of the following genotypes: *ap-Gal4, UAS-Yki, UAS-GFP, UAS-LacZ* (A) and *ap-Gal4, UAS-Yki, UAS-GFP, EP-dPGC1* (B). GFP is shown in green. DAPI labels the DNA and is shown in blue. Scale bars, 100 µm.

**(C)** Quantification of GFP-positive area in third instar wing imaginal discs of the genotypes in A and B. GFP-positive areas were normalized to the mean of the control (*ap>Yki*). Statistical significance was determined using an unpaired t-test (n = 14 [*ap>Yki*], n = 10 [*ap>Yki, dPGC1*). ns, non-significant.

**(D, E)** mRNA quantification by qPCR of *dMfn* (D) and *Opa1* (E) in the indicated genotypes. The control genotype is *ap>Yki*. Each genotype was run in five biological replicates. RP49 was used as the housekeeping gene. Statistical significance was determined using unpaired t-tests. ns, non-significant.

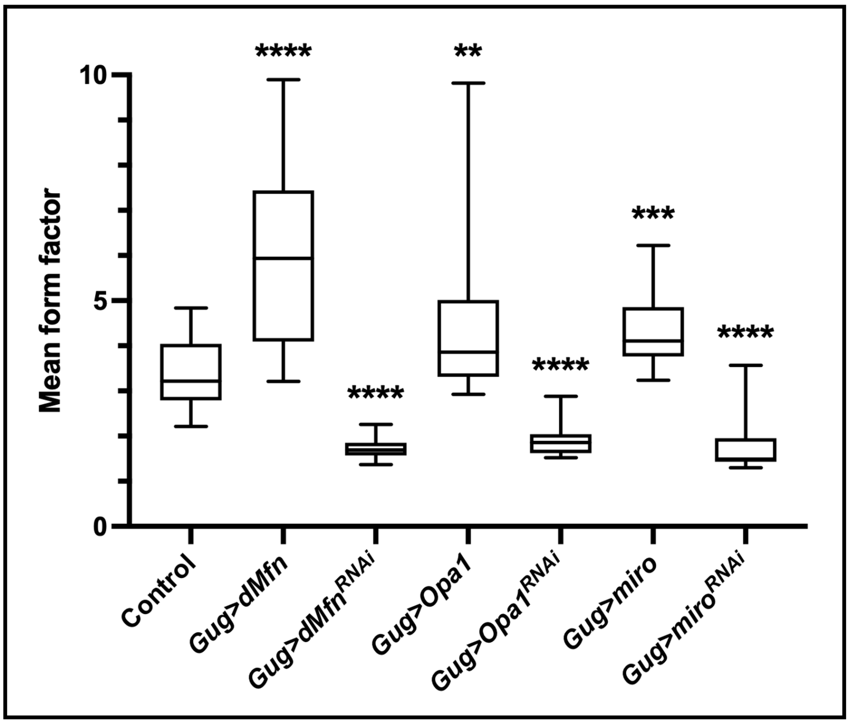


**Figure S9. Changes in Mitochondrial Shape upon *dMfn*, *Opa1*, and *miro* Gene Manipulation**

Quantification of mitochondrial mean form factor (shape measure where a value of 1 indicates a round object and values increase with elongation) from confocal micrographs obtained from the peripodial membrane of the following genotypes: *Gug-Gal4, UAS-Mito-GFP* (Control)*; Gug-Gal4, UAS-Mito-GFP, UAS-dMfn*; *Gug-Gal4, UAS-Mito-GFP, UAS-dMfn-RNAi*; *Gug-Gal4, UAS-Mito-GFP, EP-Opa1; Gug-Gal4, UAS-Mito-GFP, UAS-Opa1-RNAi; Gug-Gal4, UAS-Mito-GFP, UAS-miro*; and *Gug-Gal4, UAS-Mito-GFP, UAS-miro-RNAi.* Statistical significance was determined using unpaired t-tests with Welch's correction for parametric data and Mann-Whitney tests for non-parametric data (n = 20 [Control], n = 20 [*Gug>dMfn*], n = 20 [*Gug>dMfn-RNAi*], n = 23 [*Gug>Opa1*], n = 23 [*Gug>Opa1-RNAi*], n = 20 [*Gug>miro*], n = 20 [*Gug>miro-RNAi*]). **, $p < 0.01$, ***$p < 0.001$, ****$p < 0.0001$.

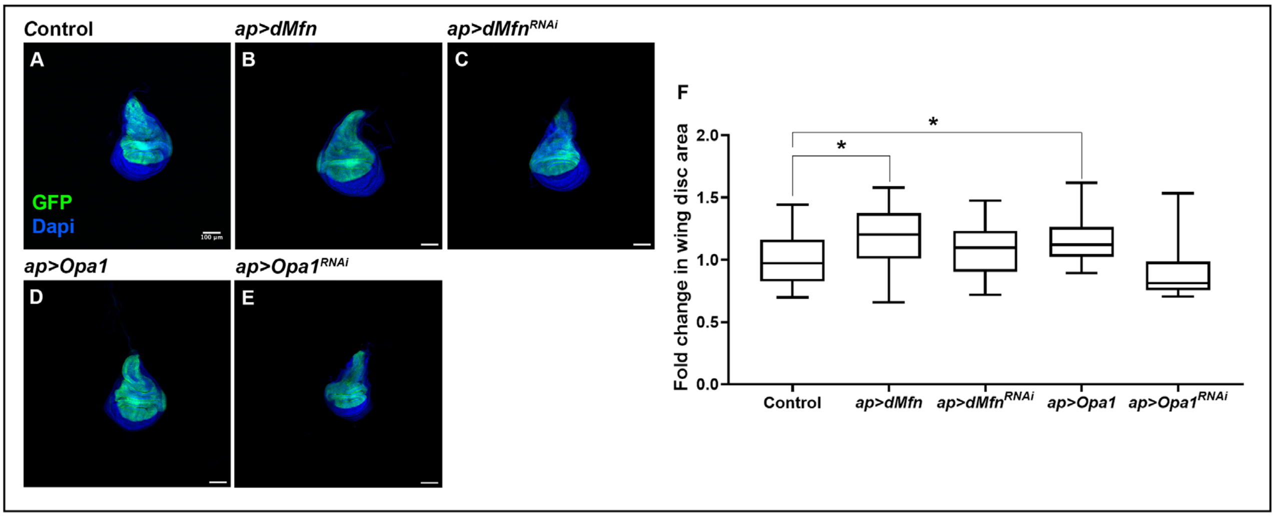


**Figure S10. Wing Disc Size upon *dMfn* and *Opa1* Manipulation**

**(A-E)** Confocal micrographs showing discs of the following genotypes: *ap-Gal4, UAS-GFP, UAS-LacZ* (A); *ap-Gal4, UAS-GFP, UAS-dMfn* (B); *ap-Gal4, UAS-GFP, UAS-dMfn-RNAi* (C); *ap-Gal4, UAS-GFP, EP-Opa1* (D); and *ap-Gal4, UAS-GFP, UAS-Opa1-RNAi* (E). GFP is shown in green. DAPI labels the DNA and is shown in blue. Scale bars, 100 µm.

**(F)** Quantification of GFP-positive area in third instar wing imaginal discs of the genotypes shown in A-E. GFP-positive area was normalized to the mean of the control (*ap>LacZ*). Statistical significance was determined using unpaired t-tests with Welch's correction for parametric data and Mann-Whitney tests for non-parametric data (n = 18 [Control], n = 18 [*ap>dMfn*], n = 20 [*ap>dMfn-RNAi*], n = 20 [*ap>Opa1*], n = 19 [*ap>Opa1-RNAi*]). *p < 0.05.

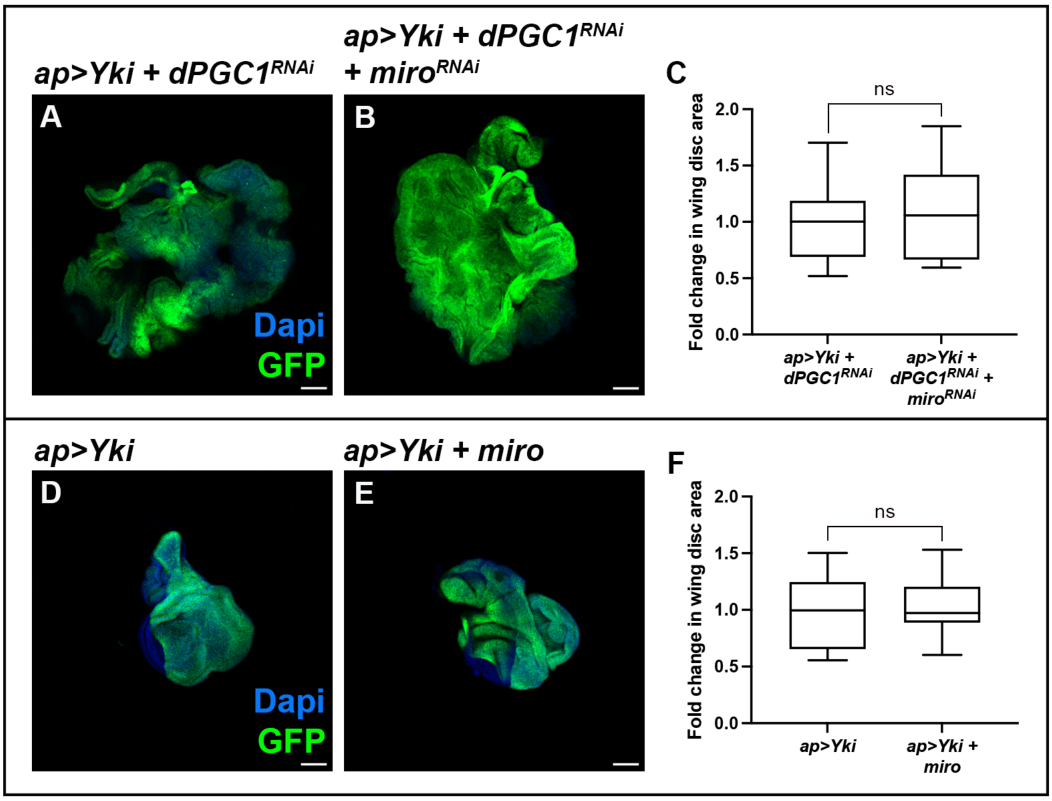


**Figure S11. Miro Does Not Affect Yki-Driven Tumors**

**(A, B)** Confocal micrographs of wing imaginal discs of the following genotypes: *ap-Gal4, UAS-Yki, UAS-GFP, UAS-dPGC1-RNAi* (A) and *ap-Gal4, UAS-Yki, UAS-GFP, UAS-dPGC1-RNAi, UAS-miro-RNAi* (B). GFP is shown in green. DAPI labels the DNA and is shown in blue. Scale bars, 100 µm.

**(C)** Quantification of GFP-positive area in wing imaginal discs of the genotypes in A and B. GFP-positive areas were normalized to the mean of the control (*ap>Yki, dPGC1-RNAi*). Statistical significance was determined using a Mann-Whitney test (n = 22 [*ap>Yki, dPGC1-RNAi*], n = 26 [*ap>Yki, dPGC1-RNAi, miro-RNAi*]). ns, non-significant.

**(D, E)** Confocal micrographs of wing imaginal discs of the following genotypes: *ap-Gal4, UAS-Yki, UAS-GFP, UAS-LacZ* (D) and *ap-Gal4, UAS-Yki, UAS-GFP, UAS-miro* (E). GFP is shown in green. DAPI labels the DNA and is shown in blue. Scale bars, 100 µm.

**(F)** Quantification of GFP-positive area in wing imaginal discs of the genotypes in D and E. GFP-positive areas were normalized to the mean of the control (*ap>Yki*). Statistical significance was determined using an unpaired t-test (n = 21 [*ap>Yki*], n = 25 [*ap>Yki, miro*]). ns, non-significant.

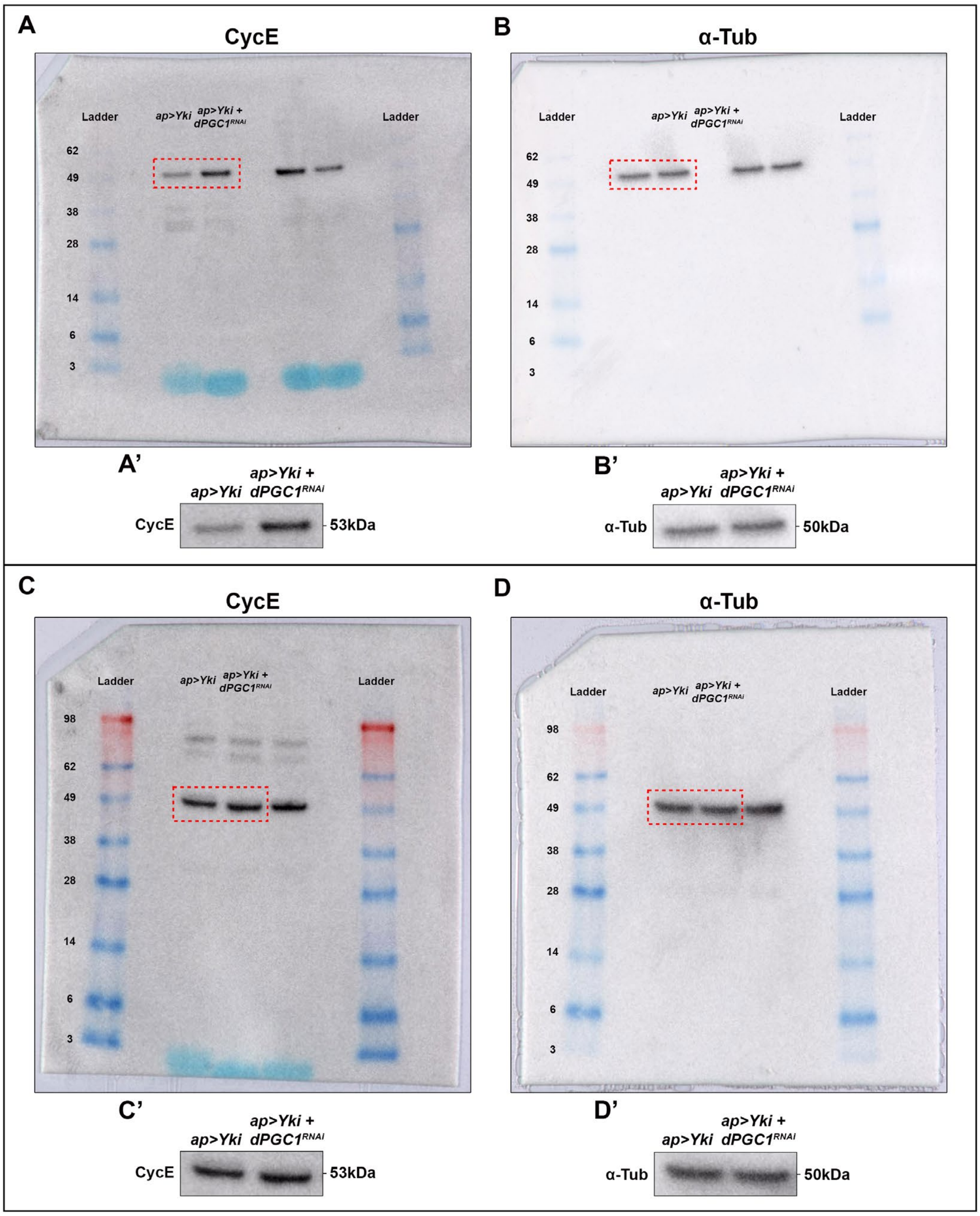


**Figure S12. Representative Whole Western Membrane Images of Cyclin E Protein Levels in *Yki* + *dPGC1-RNAi* Tumors**

**(A-D)** Examples of two western blot membranes to analyze protein levels of Cyclin E (A and C) and $\alpha$-Tubulin (B and D) in wing imaginal discs of the following genotypes: *ap-Gal4, UAS-Yki, UAS-GFP, UAS*-LacZ; and *ap-Gal4, UAS-Yki, UAS-GFP, UAS-dPGC1-RNAi.* The molecular weights (in kDa) of the visible bands of the ladder are indicated at the left of each membrane. The dashed red boxes in A-D indicate the regions of the membranes that are shown as magnifications in A'-D', respectively. Note that the western blot membrane of panels A and B corresponds to the one shown in Fig 6B and therefore panels A' and B' are the same as those in Fig 6B. The membranes shown here include additional genotypes not relevant to this study. Only the lanes corresponding to the relevant samples are indicated.

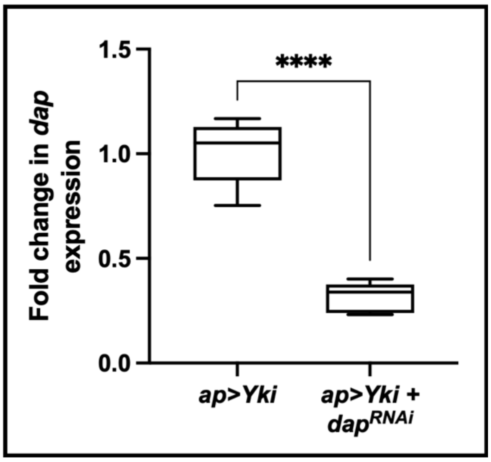


**Figure S13.** ***UAS-Dap-RNAi*** **Transgene Efficiency**

mRNA quantification by qPCR of *dap* in wing imaginal discs of the following genotypes: *ap-Gal4, UAS-Yki, UAS-GFP, UAS-LacZ* and *ap-Gal4, UAS-Yki, UAS-GFP, UAS-dap-RNAi.* The control genotype is *ap>Yki.* Each genotype was run in five biological replicates. RP49 was used as the housekeeping gene. Statistical significance was determined using an unpaired t-test. ****$p < 0.0001$.

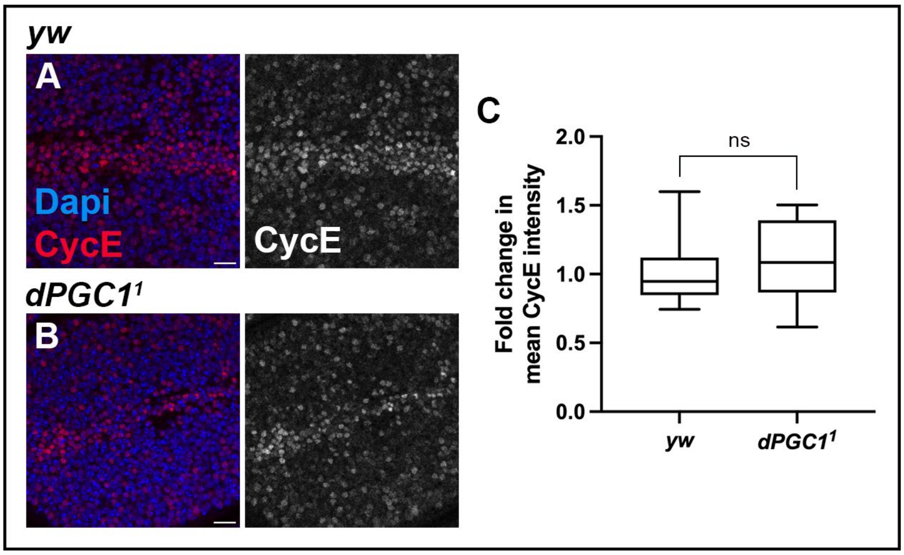


**Figure S14. Cyclin E in Wing Imaginal Discs of *dPGC1* Mutant Flies**

**(A, B)** Confocal micrographs showing the wing pouch of imaginal discs of the following genotypes: *yw/yw* (control, A) and *dPGC1*$^1$/*dPGC1*$^1$ (mutant, B). Cyclin E is shown in red. DAPI labels the DNA and is shown in blue. Scale bars, 10 µm.

**(C)** Quantification of Cyclin E protein mean intensity of the genotypes in A and B. Cyclin E intensity was normalized to the mean intensity of the control (*yw/yw*). Statistical significance was determined using an unpaired t-test (n = 22 [*yw*], n = 13 [*dPGC1*$^1$]). ns, non-significant.
